# How to use random walks for modeling the movement of wild animals

**DOI:** 10.1101/2020.03.11.986885

**Authors:** Geoffroy Berthelot, Sonia Saïd, Vincent Bansaye

## Abstract

Animal movement has been identified as a key feature in understanding animal behavior, distribution and habitat use and foraging strategies among others. At the same time, technological improvements now allow for generating large datasets of high sampled GPS data over a long period of time. However, such datasets often remain unused or used only in part due to the lack of practical models that can directly infer the desired features from raw GPS locations and the complexity of existing approaches. Some of them being disputed for their lack of rational or biological justifications in their design. We propose a simple model of individual movement with explicit parameters based on essential features of animal behavior. The main thrust was to stick to empirical observations, rather than using black-box models that could possibly perform very well while providing little insight from an ecological perspective. We used a simple model, based on a two-dimensional biased and correlated random walk with three forces related to advection, attraction and immobility of the animal. These forces can be directly estimated using individual raw GPS data. The performance of the model is assessed through 5 statistics that describe the spatial features of animal movement. We demonstrate the approach by using GPS data of 5 roe deer with high frequency sampling. We show that combining the three forces significantly improves the model performance. We also found that the model’s parameters are not affected by the sampling rate of the GPS, suggesting that our model could also be used with low frequency sampling GPS devices. Additionally, the practical design of the model was verified for detecting spatial feature abnormalities (such as voids) and for providing estimates of density and abundance of wild animals. Our results show that a simple and practicable random walk template can account for the spatial complexity of wild animals. Integrating even more additional features of animal movement, such as individuals’ interactions or environmental repellents, could help to better understand the spatial behavior of wild animals.

## Introduction

Animals live in an environment that is patchy and hierarchical, and the manner in which individuals search for spatially dispersed resources is crucial to their success in exploiting them (1). At the same time, the tracking of animals using the modern global positioning system (GPS) now allows for the collection of important datasets on animal locations (2). They are often used for the analysis of the home range behavior i.e., restrict their movements to self-limited portions of space far smaller than expected from their sole locomotion capacities (3) and, more generally, to better understand the spatial and temporal behavior of animals (4, 5). New, smaller and reliable devices allow for gathering large datasets (*e.g.* locations or activity data for instance) at a finer temporal and spatial scale and offer a greater opportunity to investigate animal movement at the individual scale. However, datasets often remain only partially used due to both the lack of practical models that can directly infer the desired features from raw GPS locations and the complexity of existing approaches. Meanwhile, ecologists in particular are called to develop new capabilities to deal with these large datasets (6, 7).

The modeling of animal movement includes a wide range of methodologies: biased and/or correlated random walks (BCR), the disputed Lévy Flight/walk (8–12), Stochastic Differential Equation (SDE) (13–16) including diffusion models based on the two-dimensional Ornstein-Uhlenbeck process (17–21) and other more exotic algorithms using *ad-hoc* rules to mimic movement features such as memory (22, 23). Lévy Flight has convenient patterns but ecological motivations are scarce (11). SDE -the continuous analog of BCR- or the Brownian bridge and Movement Model (24) may be used to interpolate the trajectory between two observations. SDE includes a drift (directional) and one or several random diffusion processes (16, 25). BCRs are convenient tools to model animal movement as the discrete time is well adapted to regular GPS data (25) and the parameters of the BCR can be directly interpreted in terms of the behavior of the animal. They correspond to the attraction of some locations, the inertia and memory feature of the movement, time dependence of the movement, local interactions with other individuals, etc. Some key features of animal movement have already been identified by previous studies, including diffusion (or randomness) which corresponds to an isotropic random motion, where the individual has the same probability to go in all directions; Attraction (directional bias) where the movement of the animal is anisotropic and is confined in an area or domain, according to (26) and other studies (3), while the attraction may depend on the distance from the isobarycenter of locations (27); Inertia where the movement of the animal is also shaped by foraging tasks where the animal alternates exploration periods -the path has high tortuosity-with straightforward movements (28). These three features can be implemented as parameters of a BCR.

This study aims at modeling animal movement of sedentary individuals over short periods (29), in a homogeneous landscape using GPS data sets and a BCR. As a first approach, we consider one individual of a given species with no interaction and simulate its movement in continuous space and discrete time in 2 dimensions using a BCR with the aforementioned parameters -diffusion, attraction, inertia- and one additional term: immobility. This late parameter takes into account the absence of movement between a pair of locations (*i.e.* distance is 0). This can be accredited to technological limitations with the satellite telemetry due to a weak GPS signal strength (*i.e.* due to natural elements: such as when the animal was standing underneath a rock or due to dense clouds, dust particles, mountains or flying objects, such as airplanes). However, this can also be part of the behavior of the animals, during specific times: sleep cycles for instance. The introduced model is general, simple and informative as the three parameters are directly inferred from the GPS data set. The model can be sophisticated by including more complicated environmental aspects of individual movement, such as spatial memory (30, 31), reinforcement and site fidelity (32), environmental predictability (33) including landscape effect (34), interacting individuals and prey-predator dynamics. We also introduce five statistics that describe the spatial features of animal movement, with a particular interest in census (*i.e.* population estimation) using transects. These statistics allow for estimating the ‘performance’ of the BCR model, or in other words its ability to mimic the spatial characteristics of an animal’s movement. Finally, we investigate how our model can address ecological questions including census and spatial issues, using GPS data sets.

## Material and Methods

### Data

The locations of 5 GPS-collared red deer (*Cervus elaphus*) were gathered at La Petite Pierre National Hunting and Wildlife Reserve (NHWR), in north-eastern of France (48.8321 (Lat.) / 7.3514 (Lon.)). The reserve is an unfenced 2670 ha forest area characteristics by deciduous trees (mostly *Fagus sylvatica*) in the western part and by coniferous species (mostly *Pinus sylvestris* and *Abies alba*) in the eastern part in nature reserve surrounded by crops and pastures. It is located at a low elevation area of the Vosges mountain range, which rises up to 400 m a. s. l. The climate is continental with cool summers and mild winters (mean January and July temperatures of 1.4 and 19.6°C, respectively, data from Phalsbourg weather station, Meteo France, from 2004 to 2017). Three ungulate species are present and mainly managed through hunting in the NHWR: wild boar, red deer and roe deer. The present study focuses on female red deer for test model. A detailed overview of the landscape and surroundings is given in (35). The GPS data had regular observation frequencies with high frequency sampling (Table 1). In the following text, we note 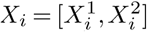 the successive locations of the individual with *X*_*i*_ ∈ ℝ ^2^ and *i* = 1, 2,…, *n*. We use *t*_*i*_ (*t*_1_ = 0) as the time elapsed between two successive locations *X*_*i*−1_ and *X*_*i*_ and

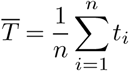

as the average sampling time. The trajectory of the animal, or ‘path’, was interpolated using linear interpolation between each pair of recorded observations (Figure 1 and detailed in Supplementary file 1 (eq. 18) and associated Graph 2). It approximates the animal travels in straight lines at constant velocity between each pair of locations (36). The home/attractor *X*_*F*_ of one individual was estimated as the isobarycenter of all recorded locations:

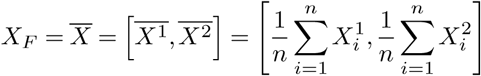

**Table 1.**
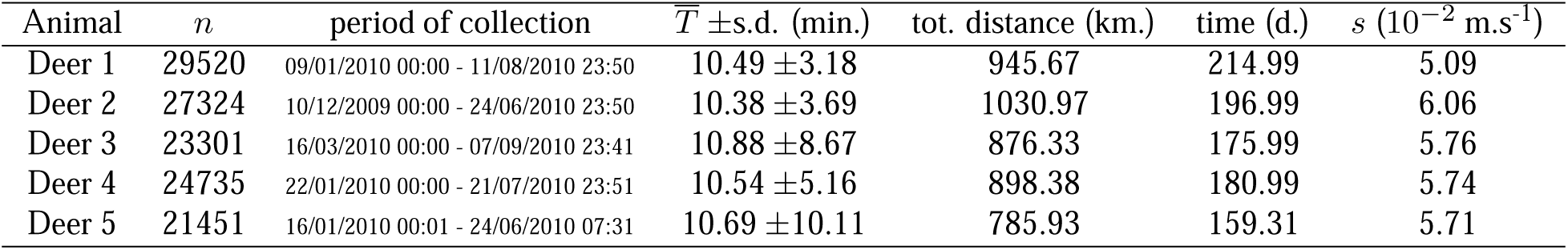
Data summary. For each animal, the total number of observations *n* is given along with the period of collection (date and time), the sampling rate 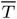 (*i.e.* the average time between 2 observations) (in min.) and corresponding standard deviation, total distance (in kilometers), total recording time (in days) and average speed *s* (in 10 ^−2^ m.s^-1^).

**Fig. 1.**
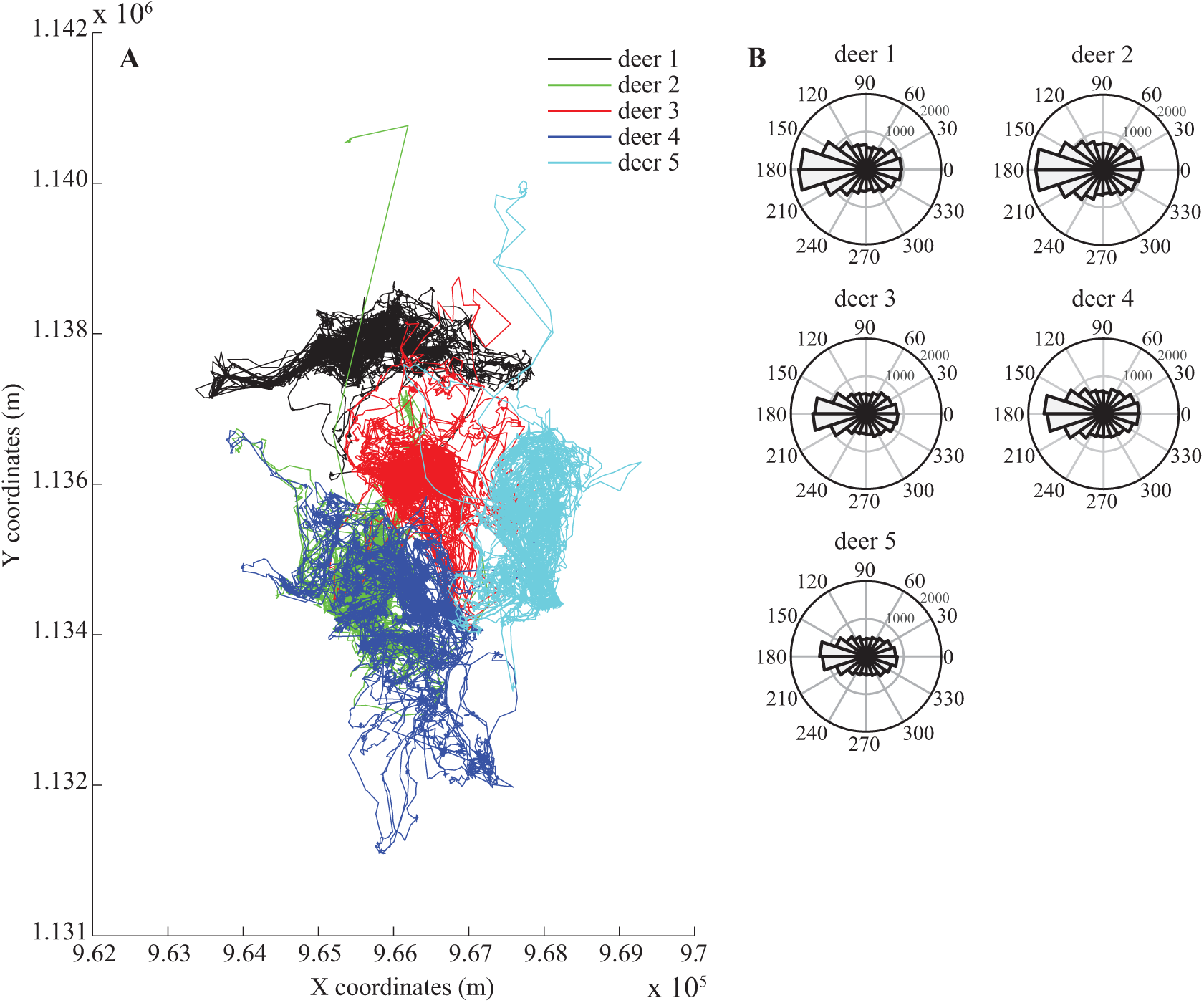
Individual paths of the five red deer. Individual paths of the five red deer. The individual paths are plotted for the five red deer (left panel, **A**) along with the distribution of the relative turning angles (degrees) in polar plots (right panel, **B**). An angular value of 0 consists in a straight motion from the previous location, while a relative turning angle of 180°C corresponds to a turn back.

### Modeling framework

The movement of an animal was modeled by a two-dimensional BCR in discrete time and continuous space which included 3 parameters coupled with isotropic diffusion:

- *Diffusion*: We considered a random direction with uniform spatial distribution in a 2D plane,
- *Attraction* (*p*_*F*_): A natural way to include this feature is to increase the probability to go to a fixed attractive point named attractor (37). This yielded a drift or advection parameter in the direction of *X*_*F*_,
- *Inertia* (*p*_*I*_): This parameter increased the probability to move forward (*i.e.* to perform one step in the direction of the previous step),
- *Immobility* (*p*_*s*_): We included this as a specific parameter and the movement was stopped for one step.

Thus, the movement of the animal was modeled by a chain *X*_*n*_, characterized by a transition matrix:

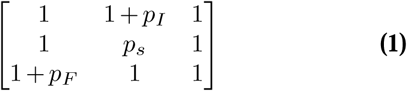

renormalized by 8 + *p*_*I*_ + *p*_*s*_ + *p*_*F*_ to obtain the probability distribution of the next position compared to the current one. In the presented case (eq. 1), the immediate past movement of the animal is coming from the down part of the matrix (*i.e.* upward vertical direction: ↑) and the attractor is estimated to be located southwest from the actual position of the individual.

We analyzed the distribution of the distances covered between each pair of locations (Supplementary file 2) and used the log-normal law to model the distance covered by the individual between each time step. Thus, if *p*_*F*_ = *p*_*I*_ = *p*_*s*_ = 0, the BCR resumes to a typical two-dimensional random walk with a log-normal step size distribution ln 𝒩(*µ,σ*^2^). The three parameters were accordingly tuned to the corresponding dataset using a straightforward estimation procedure (Supplementary file 1). We then simulated 1000 BCR and used 5 statistics that describe the spatial features of an individual’s movement to assess the BCR performance (framework detailed in Figure 2). We additionally perform a sensitivity analysis by testing the performance of the BCR but using only one or two parameters instead of the three parameters.

**Fig. 2.**
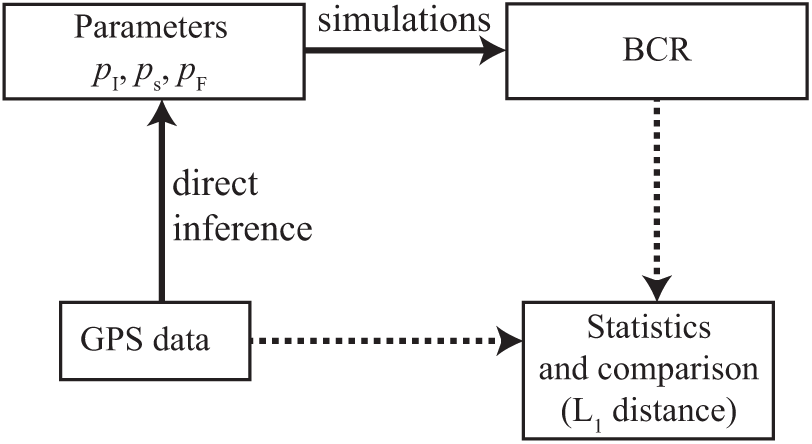
Framework used for testing the BCR model performance, for one animal. Black lines detail the two operations processed from the GPS dataset. The 3 parameters are estimated from the GPS data and -using these parameters-1000 simulations of the BCR model are computed. No particular operations are associated with the dotted black lines, but they show how the BCR and the GPS dataset are evaluated and compared using the statistics. We use the same framework to investigate the performance of the BCR with only one or two parameters.

**Fig. 3.**
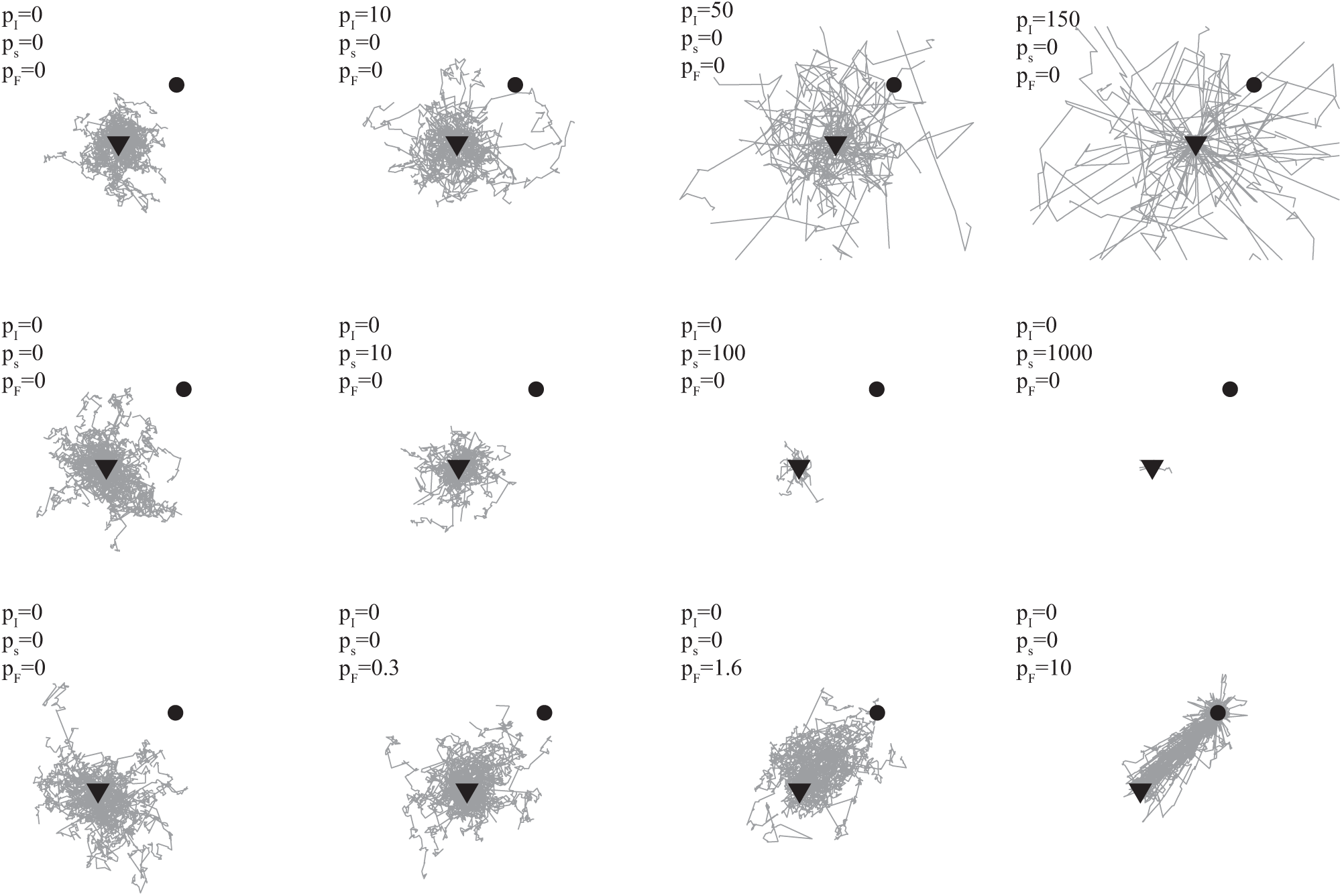
Simulated animal motions over arbitrary parameter values. Fifty motions of length *n*_*s*_ = 100 steps are simulated and originate from a common centroid (downward-pointing triangle) with increased levels of inertia (*p*_*I*_), immobility (*p*_*s*_) and attractor (*p*_*F*_). Both the location of the attractor (black dot) and the log-normal parameters controlling the step size distribution are fixed (*µ* = 3, *σ*^2^ = 1).

### Statistics for describing animal movement

The 5 statistics were designed to assess the model reliability on spatial features including: (*i*) the distribution of relative turning angles which provides information about the local movement of the animal, (*ii*) the home range which provides information about the spatial density of observations and (*iii*) observation counts using still and mobile transects, providing information on absolute observation abundance (38). We did not keep track of locations during transect counts, thus ignoring spatial information, as it was already collected in (*ii*). Additional details are provided in Supplementary file 3.

#### Distribution of turning angles

For each individual, the distribution of counter-clockwise relative turning angles 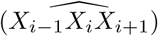 was gathered, provided *d*(*X*_*i*−1_, *X*_*i*_) *> d*_min_ and *d*(*X*_*i*_, *X*_*i*+1_) *> d*_min_ with *d*_min_ the immobility threshold distance between two locations. We set *d*_min_ *<* 10m, which corresponds to the magnitude of the error typically found in GPS locations (39). This means that we only kept the angles from observations that were separated by an Euclidean distance greater than *d*_min_.

#### Home range

We used an adaptive kernel density estimator (matlab package kde2d - kernel density estimation version 1.3.0.0) as an estimator of the utilization distribution (40) to represent the home range of the animal. The approach of Z.I. Botev provided an estimate of observation density using a bivariate (Gaussian) kernel with diagonal bandwidth matrix (41). The density was estimated over a grid of 210 × 210 nodes and we computed the home range area (in m^2^) for various values: 100, 99, 95, 90, 80, …, 20, 10% of the estimated density. Similarly to the distribution of turning angles, we compared each value of the data’s home range against the simulated one.

#### Dilation

Dilation is generally used to account for the spatial attributes of an object such as to measure an area around the path or the volume of a brownian motion (see Wiener sausage (42) and Gromov–Hausdorff distance). In our approach, we use dilation of both simulated and GPS paths for two reasons: to have a real -and comparable-number that accounts for how a trajectory has explored space and because it is natural tool from a census point of view (the dilated path corresponds to the area where the animal can be detected). Each simulated or real path was plotted in binary format in a window and dilated with a disk shape. The window size was set to a huge value in order to encapsulate the dilated path while preventing boundary effects, *i.e.* the convex envelope of the dilated area did not collide with any window border. We then estimated the surface covered by the dilated path for 100 different sizes of the disk, from disk size 1 to disk size 100. We compared each value of the data’s estimated surface against the simulated one.

#### Immobile transects

We used still transects that counted the number of times the animal was seen in their line of sight. We arbitrary set the line-of-sight value at 200m. The number of sightings of each transect was gathered and ordered in decreasing order, thus breaking the spatial dependence. We then compared the bins of the resulting histogram in the data and in the simulated path.

#### Mobile transects

First, the movement of the animal was linearly interpolated from the GPS data, meaning that between two recorded locations the individual followed a linear path. The speed of the animal between two locations was accordingly reconstructed using the recorded times *t*_*i*_ between each location. Second, we used mobile transects as the ecological sampling method, where each transect ‘count’ the intersection between its path and the animal’s one. The mobile transects followed a predefined path at a given constant speed as time increased. The area of vision of each transect was defined as a circle of a given radius. Each time the path of an individual collided with an area of vision, the count of the corresponding transect increased by 1. Two types of movements were used: linear and clockwise rotational transects. The initial locations of both types of transects are *X*_1_ and *X*_*F*_. Both the animal and mobile transects started to move at the same time. At each of the two locations *X*_1_, *X*_*F*_, 8 linear transects moved in the 8 cardinal directions, totalizing 16 transects. For the linear transects, every 10000 time steps, we set 2 × 8 new transects starting at the same locations and following the same directions. Clockwise rotational transects were rotated around *X*_1_ and *X*_*F*_ using a 500m radius. When we reached *t*_*n*_, we gathered the total count (*i.e.* the count of all transects). For the two types of transects, we gathered the total count for 6 different lines of sight: 50, 100, 200, 400, 500, 1000m. and 4 speeds: *s/*4, *s/*2, *s*, 2.*s* with *s* the average speed of the animal. We then aggregated the overall count in each of the two types of transects, and compared the results from the data and the simulated path (Supplementary file 1 and 3).

### Scale invariance

We also studied how the scale affected the BCR parameters. The movement path formed by the GPS observations *X*_*i*_ was subsampled (decimated) for each individual. We only kept every *k* observation starting with the first one and *k* ∈ [1, 10]. For *k* = 1 the path corresponded to the original one. The time spent between each successive observation was also accordingly reconstructed in order to keep track of 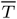 in subsampled movement paths. The time between two locations *X*_*i*_ and *X*_*i*+*k*_ was reconstructed as: 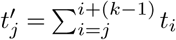 with *j* ∈ [1, 1 + *k*, 1 + 2*k*,…, *n* − (*k* − 1)].We then compared the resulting parameters *p*_*I*_, *p*_*F*_ and *p*_*s*_ as the subsampling parameter *k* increased.

### Deterministic aspects of the statistics

Whereas the BCR is a stochastic process, the deterministic aspects of the 5 statistics were tested with an increasing number of steps *n*_*s*_. The statistic associated with each realization of the model (a simulated path) is a random variable. If the distribution of these random variables has low concentration (high variance) then it is not a convenient statistic as it cannot be used as a reference for assessing the model’s performance, even when averaging over multiples realizations. On the opposite, if the statistic is deterministic it can provide a reliable tool to assess the model’s performance. This was numerically tested over a range of increasing *n*_*s*_ values with *n*_*s*_ = 10^4^, 2 × 10^4^, …, 4 × 10^5^. For each of those step values, a set of 100 BCR was simulated with parameters *p*_*I*_, *p*_*F*_ and *p*_*s*_ estimated from the first deer (see Table 2) and we studied the variance of the statistics.

**Table 2.** Estimated parameters. For each deer, the estimated parameters *p*_*I*_, *p*_*F*_, *p*_*s*_ and the two parameters *µ,σ* that control the step size distribution are given.

| Animal | $p_I$ | $p_F$ | $p_s$ | $\mu$ | $\sigma$ |
| --- | --- | --- | --- | --- | --- |
| Deer 1 | 0.01 | 0.01 | 2.01 | 2.94 | 1.01 |
| Deer 2 | 0.06 | 0.13 | 1.44 | 3.15 | 0.97 |
| Deer 3 | 0.12 | 0.05 | 1.70 | 3.07 | 1.04 |
| Deer 4 | 0.10 | 0.06 | 1.52 | 3.10 | 0.98 |
| Deer 5 | 0.22 | 0.24 | 1.66 | 3.03 | 1.06 |

### Application: detecting anomalous voids

The proposed model could be used to infer environmental and behavior information from the dataset. For example, it was possible to detect anomalous voids (or holes) in the spatial territory of the individual using Monte-Carlo simulations of the model. Anomalous means that the observed void is not related to the randomness of the movement, but rather related to a geographical artifact. The three parameters *p*_*I*_, *p*_*F*_, *p*_*s*_ and parameters of the log-normal distribution for step-size were accordingly estimated from the data of each individual, similarly to previous experiments (Figure 2 and Table 2). A simple heuristic was used to find voids in empirical and simulated paths for each individual: we computed the alpha shape of all locations using a fixed alpha radius of 60m. This allowed for determining the surface covered by all locations while preserving the voids. We then collected the area of each void provided they had an area of at least 100m^2^. We focused on voids near the center of the alpha shape in order to avoid artificial voids, generated by the weak density of locations at the boundaries. We ran 10,000 iterations of the model for each animal and estimated the probability *p*_∅_ of finding voids of different sizes in the simulated paths. This probability was then compared to voids found in the GPS datasets and available environmental information was used to determine whether any geographical element(s) could explain the unexpected voids.

## Results

The parameters estimated for each individual are given in Table 2 for the model. We showed that the parameters *µ* and *σ*^2^ were close for all individuals, and estimates of the three parameters for individuals 3 and 4 were similar. The values of *p*_*I*_ and *p*_*F*_ showed that inertia and attractor play a greater role in the movement of deer 5 (*p*_*I*_ = 0.22, *p*_*F*_ = 0.24), compared to the other individuals. The immobility *p*_*s*_ was stable across the individuals while *p*_*I*_ and *p*_*F*_ varied together (Table 2). The latter is a mechanistic effect, as they act as opposite forces. We also detailed additional configurations of the BCR where we used only some of the parameters (Supplementary files 8 and 9).

### Evaluation of the BCR model

The distribution of error of each of the model’s 5 configurations in the 5 statistics is provided in Figure 4 for deer 5 (see Supplementary file 4 for the complete results). The mean error and standard deviation, median error and interquartile range are also provided in Supplementary file 9, for all deer and statistics. We showed that combining the parameters plays an important role in modeling deer behavior. Configurations with only one parameter did not perform well on average while further investigations showed that combining *p*_*I*_, *p*_*F*_ and *p*_*s*_ allows for a better description of movement, especially regarding the census statistics for both linear and rotational transects and home range estimates (Supplementary files 4 and 9). Four of the 5 statistics became more and more deterministic and concentrated around their mean value (Supplementary file 5). On the contrary, the variance of the dilation seemed to increase for the first *n*_*s*_ steps. The trend observed in the linear mobile transects also seemed to increase, however the trend was not clear and presented small fluctuations.

**Fig. 4.**
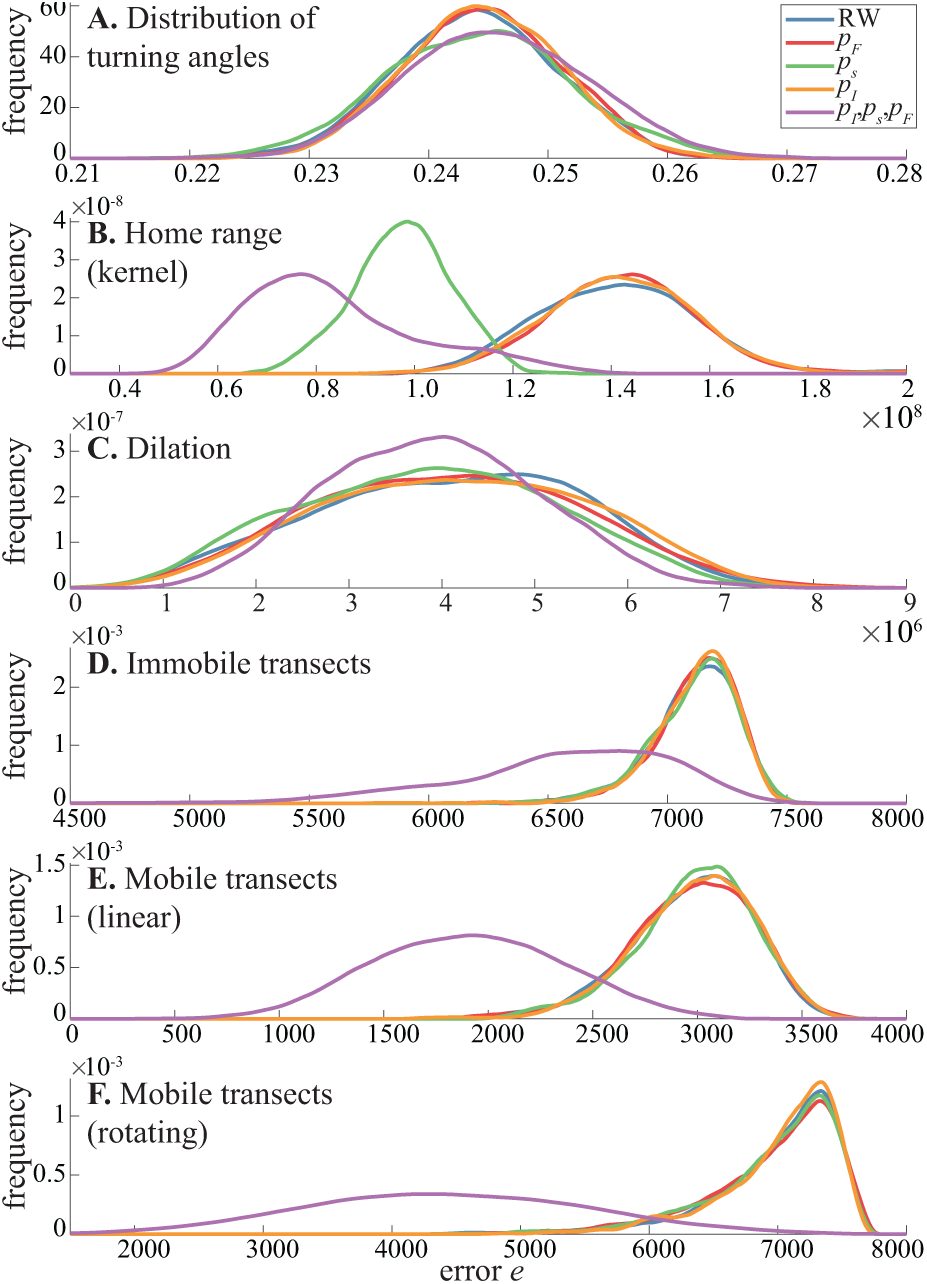
Density of error *e* of all 5 configurations tested in each statistic for deer 5. Densities are fitted by the Epanechnikov kernel function. A complete comparison of results for all red deer is provided in Supplementary file 1.

### Scale invariance

The resampling of movement paths showed that *p*_*s*_ decreased as the subsampling rate increased in all five deer. The three other parameters remain roughly constant (Supplementary file 6).

### Application: detecting spatial voids

The resulting alpha shapes and detected voids (holes) are presented for each deer in Supplementary file 7. The probability *p*_∅_ of observing such voids is computed and showed in Figure 5. Many voids whose area fell in the interval [0, 1.5 × 10^4^] were related to boundary conditions, where the alpha shape produced artificial voids due to less dense areas. However, the Monte Carlo simulations show that 3 voids, located inside the alpha shape (void 1 (deer 1) and voids 1 and 2 (deer 4)), should not appear. In other words, these voids are possibly not related to movement randomness but to other spatial features, with good probability. The distributions of errors for each configuration varied in each deer.

**Fig. 5.**
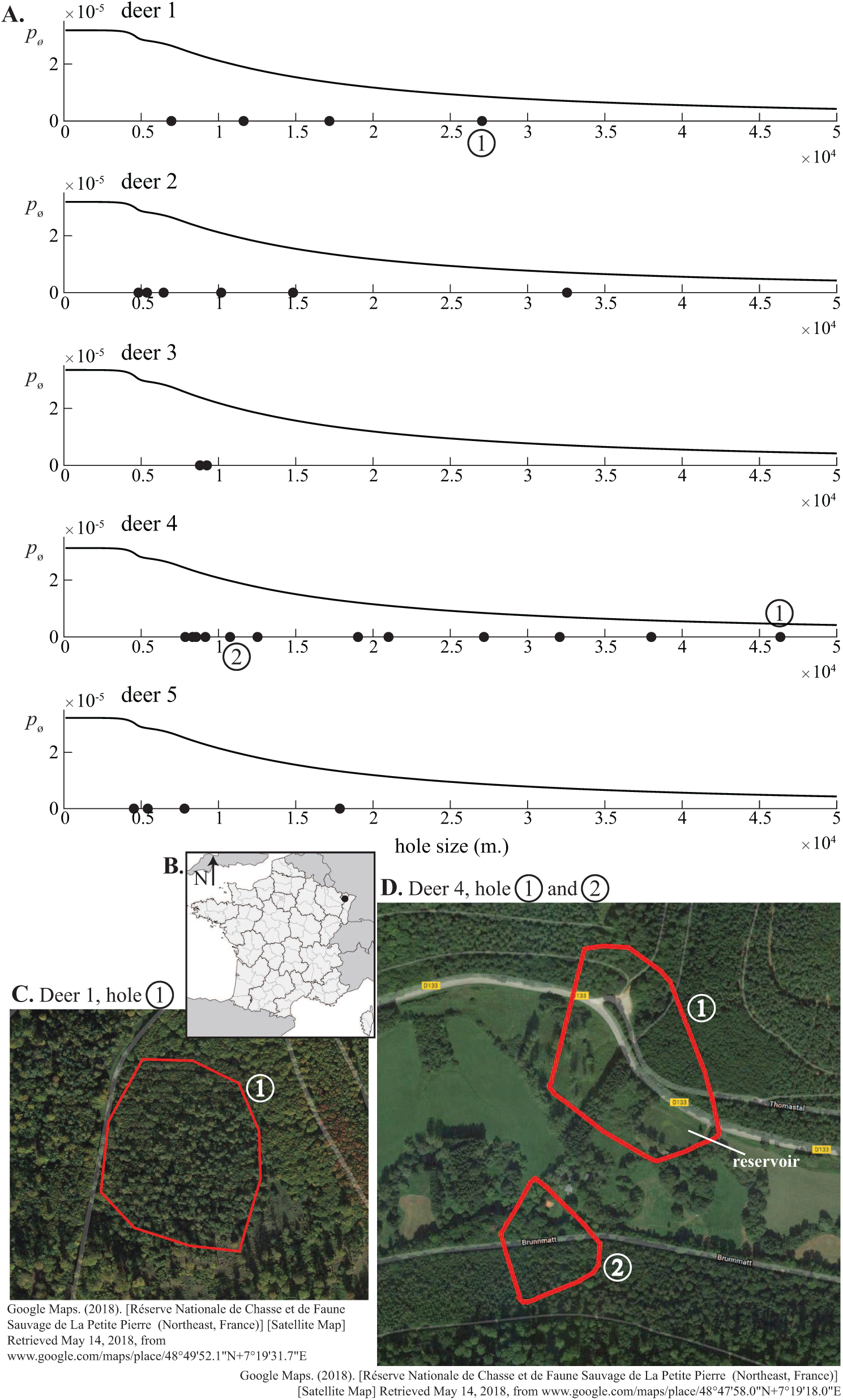
Using the framework and model to identify spatial voids in movement paths. In panel **A** the probability *p*_∅_ of finding voids of different sizes in the simulated paths is given for each deer (black lines). All voids *>* 100m^2^ detected in the empirical GPS location are given (black dots in x-axis). Selected voids (circled numbers) correspond to voids that both are near the center of the alpha shape and have a low *p*_∅_ (see alpha shape figures in Supplementary file 1). **B** Geographic location of the study area (black dot). The three voids detected in deer 1 and 4 are detailed in panel **C** and **D** along with the environmental features. Image in **B** was created by TomKr and is distributed under GNU Free Documentation License. It corresponds to the map of France with regions and departments in equirectangular projection and was realized with free IGN data base GeoFla (www.ign.fr).

## Discussion

In this work we aim at providing valuable and reliable ecological information regarding the components of animal movement. We introduce a simple and tractable model to deal with animal movement, based on a two-dimensional BCR in discrete time and continuous space that allows for combining the ecological forces in a simple way. The parameters of the BCR are directly estimated using the GPS data recorded in a large herbivore and its performance is assessed in 5 spatial and ecologically-related statistics. Four of them differ from the typical signals or parameters calculated based on empirical relocations (43) and address the home range size, census issues and animal behavior. The framework is presented in Figure 2 and explains how synthetic (or simulated) paths parameterized from empirical observations are ultimately compared to the empirical paths throughout the statistics. It describes how any field data (30), whatever the sampling rate and type, should be analyzed whenever possible. However, we emphasize that the attributes of the data (such as the sampling rate for instance) should be consistent amongst different individuals in order to allow for inter-individual comparison (see (44) and subsequent discussion). It is also important to have a sufficient number of locations (‘sample size’) as precision in parameter estimation scales with the sample size, meaning that the more observations, the higher the precision. We focus on three essential forces that may allow for an efficient description of animal motion over large periods of time: inertia *p*_*I*_, immobility *p*_*s*_ and attractor *p*_*F*_. Other perturbations are encapsulated into a random noise. The BCR is then designed to embed the main ecological features driving the movement of most terrestrial animals: exploratory or foraging behavior (inertia), patch exploitation or sleeping periods (immobility), attractor (reference and working-memory). The results display that by using those parameters, we get a much better description of animal movement compared to an unbiased random-walk with a log-normal step size distribution.

The model reproduces the distribution of relative turn angles observed in a large herbivore, provided the parameters are tuned accordingly. This distribution of turning angles is similar in all five individuals and resembles a *π*-oriented oval (Figure 1). This oval pattern was already noted in many other species such as the elk (45), the brushtail possum (‘possum’: *Trichosurus vulpecula*) (44), the caribou (46), the Mediterranean mouflon (*Ovis gmelini musimon Ovis*) (47) and even in Cinereous Antshrike (*Thamnomanes caesius*) flocks (48). GPS errors are known to possibly generate such directional bias where a stationary animal is most likely to be measured as turning 180°C or moving towards (49). We constructed turning angles using locations separated by at least *d*_min_ and this pattern remains and carry additional analyses using the adehabitatHR package in R (50) to *i* compute the residence time as the time spent into a 100m. circular area for each localization and to *ii* study the relative turn angles associated with high (*>* 100m.) or small (*<* 100m.) distances covered during a movement. We find that frequent turn backs are associated with high residence time. We also find that turn angles are more evenly distributed in the small distances and that small turn angles are more associated with small distances values. These analyses suggest that the animals have an exploratory / foraging behavior in a local patch. Exploratory or foraging behavior is often modeled using stochastic processes satisfying the Markov property, such as the bivariate Ornstein-Uhlenbeck diffusion process which produces similar distribution patterns of relative turn angles (51, 52), in line with our BCR approach.

Comparison of model configurations reveals that both the two-dimensional random walk and the configurations with only one parameter do not perform well in most cases. On the contrary, combining all three parameters *p*_*I*_ (inertia), *p*_*s*_ (immobility) and *p*_*F*_ (attractor) provides better results in the vast majority of cases (Figure 4 and Supplementary files 4 and 9). This confirms that animal movement is a complex process, driven by several forces instead of a single and dominant one.

The inertia, describing the short-term memory effect, is the first force introduced in this approach. Whether the use of land space by the animal is dependent on short-term or longterm memory is a debated topic. It gave rise to a series of studies that emphasized the importance of memory in animal movement from a biological or modeling perspective (30, 31, 53–57). These studies also underlined that inferring memory effects directly from relocations is not a trivial task. Those relocations instead depend on a mixture of effects, including landscape and territorial constraints, resource patches and possibly long-term memory. Using a single memory feature *p*_*I*_ might be a too simple approximation for efficiently capturing the memory effect. In our approach it is possible to alter *p*_*I*_ in order to include several previous steps instead of just one.

Immobility combines several features of animal movement including animal at rest, in vigilant state, and GPS noise. Multiple factors are known to affect GPS noise, including topographic exposure, canopy cover, vegetation height and the slow movement of the ionosphere. The latter changes by a few centimeters during 30sec intervals (2), possibly introducing up to 20 fold this bias in each of the recorded GPS observations. However, this is small regarding the average step size of non-immobile movements, ranging from 41m (red deer 1) to 46m (red deer 3 and 5). Thus, we assume that the measured step lengths and turning angles reflect the reality. Immobile (*i.e.* ≤ *d*_min_) observations represent a large proportion in our total datasets: 25.0% (red deer 1), 17.2% (red deer 2), 22.0% (red deer 3), 19.0% (red deer 4) and 23.6% (red deer 5), associated with specific behaviors such as on-site foraging, eating, resting, etc. The estimates of *p*_*s*_ in all five animals are greater than inertia or attractor (Table 2), underlining the importance of considering immobility when analyzing the movement of red deer. This is in line with previous experimental studies that showed the high frequency of feeding, resting cycles in red deer and labile diet (58).

Site fidelity is the recurrent visit of an animal to a previously occupied location. This is a well-known and wide-spread behavior in the animal kingdom (59). The animal favors locations that are ecologically valuable and related to a foraging or explorative behavior. In our approach, we rather and simply depict site fidelity using one single attractor *p*_*F*_. The fact it improves the performance of the model when combined with inertia and immobilism confirms that site fidelity (or a simplified estimation of it) should be taken into account when modeling deer movement.

Other analyzes were carried out to ensure the robustness and consistency of the BCR model, including the impact of the GPS sampling rate on the estimated parameters. Several authors pointed out that the temporal resolution of the discretization is of importance: it should be relevant to the considered behavioral mechanisms (5, 44, 60, 61). Schlägel and Lewis focused on the quantification of movement models’ robustness under subsampled movement paths (61). They found that increased subsampling leads to a strong deviation of the central parameter in resource selection models (61, 62). They also underlined that the parameter estimates vary with sampling rate when movement models are fitted to data. Postlethwaite and Dennis highlighted the difficulty of comparing model results amongst tracking-datasets that vary substantially in temporal grain (44)). We use data with a relatively high sampling rate (roughly 10m.) and a period of study that is appropriate to the analysis of animal movement at the year scale (Table 1). More importantly, the five animals have the same sampling rate (Table 1). We changed the sampling rate of the movement path to ensure that the parameters related to directional movement are scale invariant (Supplementary file 6). We found that they remain almost constant with increased subsampling, thus strengthening their essential role in animal movement (Supplementary file 6). On the other hand, distance-related parameters such as *p*_*s*_, *µ,σ*^2^ are highly sensitive to the subsampling rate, but it is a mechanistic effect of the subsampling procedure: as we progressively prune the *i*^*th*^ values, the distance between each GPS location increases. The distribution of turning angles or the home range estimates can vary with the temporal scale of study and sampling rate of datasets (44, 63). Schlägel and Lewis further underlined that important quantities derived from empirical data (*e.g.* travel distance or sinuosity) can differ based on the temporal resolution of the data (61, 62). This could have introduced bias in the aforementioned statistics and in parameter inference. However, both the sampling rate and temporal scale of study are similar in all our datasets (Table 1), thus allowing for unbiased inference, proper evaluation of the model’s performance and inter-individual comparison. We also investigated the deterministic feature of the 5 statistics and found that the variance decreases or does not change as the number of simulated steps increases in most of the statistics. Thus, 4 of the 5 statistics are robust and add limited randomness to the results when the number of steps increases. The long-term trend is not clear in the mobile transects case as we investigated the variance over 4 × 10^5^ steps and we may only observe a transient increase or stagnation. This statistic is expected to be similar to the one of immobile transects but the speed of convergence to the null variance may be very slow and it may take a much larger number of steps. The variance of the estimated areas in the dilation statistic increases with *n*_*s*_ because we dilated the simulated paths in a huge window, encapsulating the whole path including a very large portion of empty space around it. This was done to prevent boundary effects when assessing the area of dilated paths, *i.e.* to make sure that dilated paths do not hit any of the window bounds. Otherwise this would produce biased, underestimated areas. However, using a smaller window or, again, a much larger number of steps would result in a null-variance.

Another leading rational of this work is to investigate the model’s ability to address ecological challenges such as estimating the abundance of a given species (census) or detecting anomalous spatial features. We use both mobile and immobile simulated transects to illustrate how the model could be used in the first problem. The probability of counting the same animal multiple times can then be estimated using Monte Carlo simulations. The second issue is, for instance, to detect spatial voids in empirical movement paths. The location of animals may present empty spatial voids (or holes) of various sizes. This may be related to environmental conditions such as urban areas, water, cliff, or other ecological reasons (such as interactions with other individuals e.g. repulsive marks) and other factors. Using numerical simulations of the model, we are able to detect anomalous voids in the dataset, that are not related to randomness but to human activity. The void 1 (deer 1) reveals that some environmental changes took place between the recording time of the GPS location in 2010 and the satellite image in 2018. After cross-checking with additional information from the OFB, we learned that the identified area was a forest enclosure. This explains why the deer was not able to reach this area. Both voids 1 and 2 of deer 4 also are related to human activities: forest roads, buildings and one artificial reservoir.

## Conclusions

This work introduces a tractable model, based on a two-dimensional BCR for describing animal movement in discrete time and continuous space. The model allows for a direct and explicit estimation of the three parameters that provide the optimal design regarding the studied statistics. Moreover, it allows for deriving reliable (*i.e.* independent from the GPS sampling rate) and quantitative information about the components of animal movement. Results show that combining the parameters is a key component in modeling movement, while allowing for an accurate description of the turning angles, home range size and census issues. The model also has practical applications for addressing ecological issues such as census or spatial anomalies. While we only focus on 5 animals to demonstrate the approach, the model is general and can be applied to any other species. We considered only one attractor per animal in the proposed approach and both the existence and influence of multiple attractors are yet to be investigated. Additional behavior features such as the spatial reinforcement, memory of *n* previous steps, activity rhythms (such as the circadian cycle), distance from the attractor, landscape/habitat effect (34), interactions with other animals and topological issues are currently being investigated and will be included in a future work. Another point of interest is the development of a continuous version of the proposed model, where the direction of the step is drawn from a specific distribution, whose parameters are yet to be empirically characterized.

## ACKNOWLEDGEMENTS

We are very grateful to the French Biodiversity Office (Office Français de la Biodiversité) for the data. We also thank Sylvain Billard, Jean-Michel Gaillard and Clément Calenge for the discussions and Catherine Carter for her review of the manuscript. This work was supported by the Chair ‘Modélisation Mathématique et Biodiversité’ of Veolia Environnement-Ecole Polytechnique-Museum National d’Histoire Naturelle-Fondation X and by the ANR ABIM 26(ANR-16-CE40-0001) and ANR Mov-It (ANR-16-CE02-0010).

## Ethics approval and consent to participate

All procedures performed in studies involving human participants were in accordance with the ethical standards of the institutional and/or national research committee and with the 1964 Helsinki declaration and its later amendments or comparable ethical standards. All applicable institutional and/or national guidelines for the care and use of animals were followed. The research program is hosted by the French National Hunting and Wildlife Agency (Office National de la Chasse et de la Faune Sauvage). This institution has granted all consents necessary for the fieldwork. Game captures were conducted in accordance with European and French laws. The experiment was designed to minimize animal stress and handling time and to ensure animal welfare, as defined in guidelines for the ethical use of animals in research. A specific accreditation was also delivered for capturing animals for scientific and wildlife management purposes. Animal captures and experimental procedures were in line with the French Environmental Code (Art. R421-15 to 421-31 and R422-92 to 422-94-1).

