## Supplementary file 1 - Materials and Methods for "How to use random walks for modeling the movement of wild animals"

### Supplementary file 1 - Material and Methods

We here provide additional information regarding some technical aspects of the BCR. We also further detail the statistics used.

### 1 Model design

We first describe the procedure used to estimate the parameters (see framework, Fig. 2) then further explain the configurations used. The main issue of the model is to estimate the new location at the next time step, given the actual location $X$ at time $t$:

$$X_{t+1} = f\left(X_t\right)$$

such that the function $f(\cdot)$ is assumed to be representative of the behavior of the animal on sufficiently large time scales. In our approach, the motion at a given time $t$ is represented by matrix (1) but the individual's space of directions is a continuous compact space: exact directions are used to determine the motion toward den or inertia. In the following text, we use indices $i$ instead of $t$ to underline that we simulate a sequence of steps rather than a time.

#### 1.1 Parameters

We take into account three main features of animal motion: Inertia $I$, immobilism $s$ and den attraction $F$. Considering the past $(X_{i-1})$, current $(X_i)$ and future $(X_{i+1})$ locations:

$$I := \{i : -\pi + \pi/8 < (\widehat{X_{i-1}X_iX_{i+1}}) \leq \pi + \pi/8\} \tag{3}$$

$$s := \{i : d\left(X_i, X_{i+1}\right) \leq 10\} \tag{4}$$

$$F := \{i :\mid \widehat{X_{i-1}X_tX_{i+1}} - \widehat{X_{i-1}X_iX_F} \mid\leq \pi + \pi/8\} \tag{5}$$

The minimal distance to cover is $d_{\min} \leq 10$m, and is designed to encapsulate GPS error and foraging behavior.

#### 1.2 Parameter estimation

Let a state be the 2-tuple containing the actual and previous observation $\{X_{i-1}X_i\}$. We define the two 'conflicting' states when the animal may possibly be in two states at once:

$$\mathcal{H}_{IF} := \{i : \widehat{X_{i-1}X_iX_F} \leq 0 \pm \pi/8\} \tag{6}$$

when the animal is already heading toward the den $X_F$, and:

$$\mathcal{H}_{Is} := \{i : d\left(X_{i-1}, X_i\right) \leq d_{\min}\} \tag{7}$$

when the distance between two consecutive observations is small ($d_{\min} \leq 10$ m.), describing an individual that is already immobile. The notation $d(a, b)$ refers to the Euclidean distance between two observations $a$ and $b$. The subset of non-conflicting states is:

$$\mathcal{H} := \{1, \cdots, n\} - \mathcal{H}_{IF} - \mathcal{H}_{Is} \tag{8}$$

We now define a 'triplet' $\mathcal{T}$ as the 3-tuple containing the actual, previous and future observation $\{X_{i-1}X_iX_{i+1}\}$. The subset $\mathcal{T}$ denotes a triplet in $\mathcal{H}$, while the two subsets $\mathcal{T}_{Is}$ and $\mathcal{T}_{IF}$ correspond to the triplets in the conflicting states $\mathcal{H}_{Is}$ and $\mathcal{H}_{IF}$ respectively. Estimates of the 3 parameters $p_I$, $p_s$, $p_F$ are then performed. We do not use immobile locations (i.e. distances separating two observations must be $> d_{\min}$) for these estimations. We define $x_1$, $x_2$ and $x_3$ as:

$$\begin{cases} \hat{x}_1 = \dfrac{\#\mathcal{T}_I \cap \mathcal{H}}{\#\mathcal{H}}; & x_1 := \dfrac{1 + p_I}{\chi} \\[2ex] \hat{x}_2 = \dfrac{\#\mathcal{T}_s \cap \mathcal{H}}{\#\mathcal{H}}; & x_2 := \dfrac{p_s}{\chi} \\[2ex] \hat{x}_3 = \dfrac{\#\mathcal{T}_F \cap \mathcal{H}}{\#\mathcal{H}}; & x_3 := \dfrac{1 + p_F}{\chi} \end{cases} \tag{9}$$

Let $\chi$ be the sum of matrix (1): $\chi = 8 + p_I + p_s + p_F$. The values of $x_1$, $x_2$ and $x_3$ are computed for each animal's motion. Solving eq. 9 for $\chi$ yields:

$$\chi = \frac{6}{1 - (x_1 + x_2 + x_3)} \tag{10}$$

Replacing in eq. 9:

$$\begin{cases} p_I = x_1\chi - 1 \\ p_s = x_2\chi \\ p_F = x_3\chi - 1 \end{cases} \tag{11}$$

We assume that $p_{IF} = p_I + p_F$ in $\mathcal{H}_{IF}$ and $p_{Is} = p_I + p_s$ in $\mathcal{H}_{Is}$ as a convenient arrangement and ignoring higher order conflicting cases. The same estimation procedure is used for other configurations and quantity $\chi$ is accordingly calculated depending on the number of parameters used.

##### 1.3 Standard model

The standard model contains all three parameters: $p_I$, $p_s$ and $p_F$ for describing animal motion (refers to the transition matrix (1). It offers a trade-off between the number of parameters and the description of animal motion. Let $M_0$ be the transition matrix given in eq. (1), such that at a given location, this transition matrix is:

$$M_0 = \begin{bmatrix} 1 & 1 + p_I & 1 \\ 1 & p_s & 1 \\ 1 + p_F & 1 & 1 \end{bmatrix}$$

provided each force is distinct one another ($\mathcal{H}$ state), assuming the den is located in the lower left direction and that the previous motion was oriented in the top direction. Conflict between the forces may occur (see eq. 6, 7), when the individual is performing consecutive steps toward the den (ie. in $\mathcal{H}_{IF}$) for instance:

$$M_0 = \begin{bmatrix} 1 & 1 + p_I + p_F & 1 \\ 1 & p_s & 1 \\ 1 & 1 & 1 \end{bmatrix}$$

or when the individual is considered immobile for successive steps (ie. in $\mathcal{H}_{Is}$):

$$M_0 = \begin{bmatrix} 1 & 1 & 1 \\ 1 & p_s + p_I & 1 \\ 1 + p_F & 1 & 1 \end{bmatrix}$$

When simulating a step in the model, the motion in $\mathcal{H}$ is described by:

$$f(X_i) = \begin{cases} \{X_i^1 + d\cos(\alpha_1); X_i^2 + d\sin(\alpha_1)\} & \text{if } x \in [0, 8[ \\ \{X_i^1 + d\cos(\alpha_2); X_i^2 + d\sin(\alpha_2)\} & \text{if } x \in [8, 8 + p_I[ \\ X_i & \text{if } x \in [8 + p_I, 8 + p_I + p_s[ \\ \{X_i^1 + d\cos(\alpha_3); X_i^2 + d\sin(\alpha_3)\} & \text{else} \end{cases} \tag{12}$$

with $x \sim \mathcal{U} \in [0, \chi]$, $d \sim \ln\mathcal{N}(\mu, \sigma)$, $\alpha_1 \sim \mathcal{U} \in [0, 2\pi]$, $\alpha_2 = \tan^{-1}(X_{i-1}, X_i)$, $\alpha_3 = \tan^{-1}(X_F - X_i)$ and $\tan^{-1}$ is the arctangent function.

The motion in $\mathcal{H}_{Is}$ is:

$$f(X_i) = \begin{cases} \{X_i^1 + d\cos(\alpha_1); X_i^2 + d\sin(\alpha_1)\} & \text{if } x \in [0, 8[ \\ X_i & \text{if } x \in [8, 8 + p_I + p_s[ \\ \{X_i^1 + d\cos(\alpha_3); X_i^2 + d\sin(\alpha_3)\} & \text{else} \end{cases} \tag{13}$$

The motion in $\mathcal{H}_{IF}$ is:

$$f(X_t) = \begin{cases} \{X_t^1 + d\cos(\alpha_1); X_t^2 + d\sin(\alpha_1)\} & \text{if } x \in [0, 8[ \\ X_t & \text{if } x \in [8, 8 + p_s[ \\ \{X_t^1 + d\cos(\alpha_2); X_t^2 + d\sin(\alpha_2)\} & \text{else} \end{cases} \tag{14}$$

#### 1.4 Investigation of other configurations

We also studied the performance of all the configurations of the BCR, using no parameters at all (two dimensional random walk), only one or two parameters. As an example, we define the two dimensional with log-normal step size distribution (no parameters) as:

$$f(X_i) = \left\{X_i^1 + d\cos(\alpha); X_i^2 + d\sin(\alpha)\right\} \tag{15}$$

with $d \sim \ln\mathcal{N}(\mu,\sigma)$, $\alpha \sim \mathcal{U} \in [0, 2\pi]$.

### 2 Statistics

In order to assess the BCR performance in each statistic, we use the $L^1$ norm to compare the differences between the statistic $\widetilde{\mathcal{S}}$, computed over a simulated path and the statistic $\mathcal{S}$, computed over the data-set:

$$e := \sum \text{errors} = \sum_{k=1}^{N} |\mathcal{S} - \widetilde{\mathcal{S}}_k\ | \tag{16}$$

where $e$ is the estimated difference (or error) in the given statistic, and $k = 1, \ldots, N$ the number of simulations of the BCR.

#### 2.1 Distribution of turning angles

We focus on the normalized histogram $m_j$ containing $j = 1, \ldots, 20$ bins in the $[-\pi, \pi]$ interval. The function $m$ counts the number of angular values that fall into each of the bins. The normalized histogram $m_j$ meets the following condition:

$$\sum_{j=1}^{20} m_j = 1$$

This normalization remove the effect of immobilism $p_s$ as $m_j$ is only based on non-immobile ($d > d_{\min}$) observations. We rather focus on the shape of the distribution than on the absolute number of values in each bin. The differences $S - \widetilde{S}$ between the data and the simulated paths in each of the 20 bins are computed as:

$$|\mathcal{S} - \widetilde{\mathcal{S}}_k\ | = \sum_{j=1}^{20} \left| \frac{m_j}{\sum_{j=1}^{20} m_j} - \frac{m_{j,k}}{\sum_{j=1}^{20} m_{j,k}} \right|$$

where $m_{j,k}$ is the number of angular values in bins $j$ for iteration $k$ of a given realization.

#### 2.2 Immobile transects

A mesh $m$ is defined with $r$ nodes that encapsulate all the locations $X_i$ of the animal (supporting Fig. 2 and graphic 1):

$$\begin{aligned} x_0 &= \min(X_i^1) \\ y_0 &= \min(X_i^2) \\ x_M &= \max(X_i^1) \\ y_M &= \max(X_i^2) \end{aligned} \tag{17}$$

$[x_0, y_0]$ and $[x_M, y_M]$ being the boundaries of $m$ with a constant spacing value $a$ between each node. Transects counts are then gathered and ordered in increasing order. We previously compared several meshes with different resolutions (ie. different values of $a, r$) and it only impacted the results by a scale factor.

#### 2.3 Mobile transects

The issue is to find the location of both the animal and the mobiles transects at each time step.

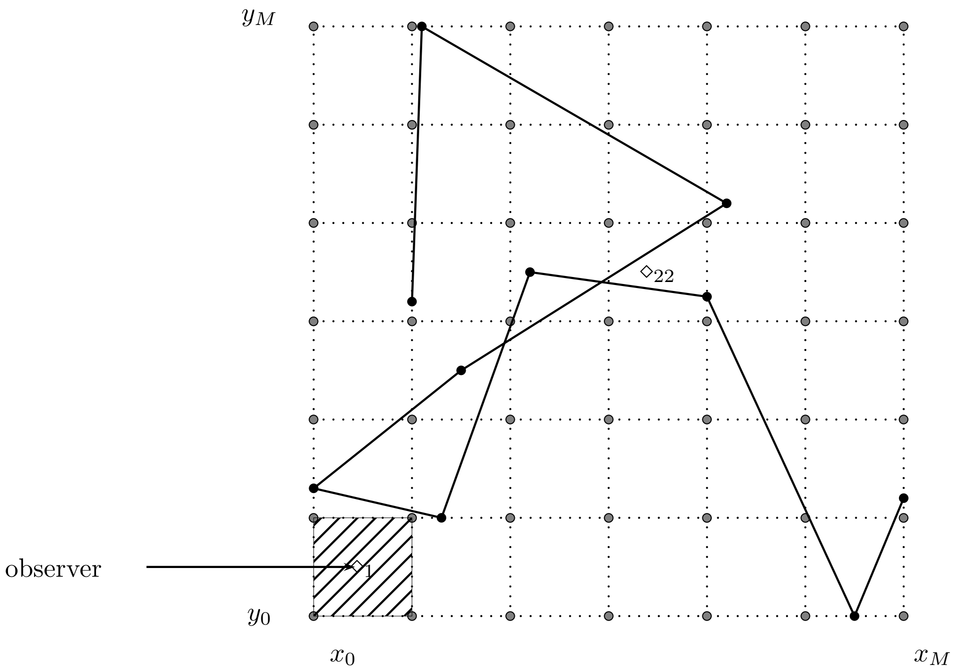


Graphic 1 – A $7 \times 7$ mesh ($r = 49$ nodes) of 36 immobile transects. Each cell of the mesh corresponds to an observation area of one immobile transect. The shaded area of the lower left cell corresponds to the area of vision of the transect $\diamond_1$. Transects are ordered in a column zig-zag way. The path (black line and dots) is counted in each cell, such that the value of $\diamond_1$ is 0 and $\diamond_{22} = 2$.

###### 2.3.1 Location of the animal

The path of the animal is reconstructed from the locations $X_i$. Linear interpolation is used between each pair of recorded locations, as detailed in Lonergan et al. (2009). It assumes the animal travels in straight lines at constant velocity between each pair of locations. Such that the path of the animal is a piecewise linear function of the observations:

$$X_i^2 = \begin{cases} \alpha_1 \cdot X_1^1 + \beta_1 & \text{if } t \leq t_{X_2} \\ \alpha_2 \cdot X_2^1 + \beta_2 & \text{if } t_{X_1} < t \leq t_{X_3} \\ \quad ... & \\ \alpha_n \cdot X_n^1 + \beta_n & \text{if } t_{X_{n-1}} < t \leq t_{X_n} \end{cases} \tag{18}$$

such that, for any time $t$ we can find the corresponding observation $X_i$ and define the segment $[X_i, X_{i+1}]$ where the animal is located (graph. 2). Thus the corresponding location of the animal is:

$$X_t = \lambda X_{i+1} + (1 - \lambda) X_i \tag{19}$$

with:

$$\lambda = \frac{t - t_{X_i}}{t_{X_{i+1}} - t_{X_i}} \tag{20}$$

###### 2.3.2 Location of the mobile linear transect

Let $s$ be the speed of transects and $\delta$ the time step. The path of a linear transect with time is:

$$X_{t+\delta} = \begin{cases} X_{t+\delta}^1 = X_t^1 \pm q \cdot s \cdot \delta \\ X_{t+\delta}^2 = X_t^2 \pm q \cdot s \cdot \delta \end{cases} \tag{21}$$

where $q = \{0, 1\}$ depending on vertical or horizontal paths.

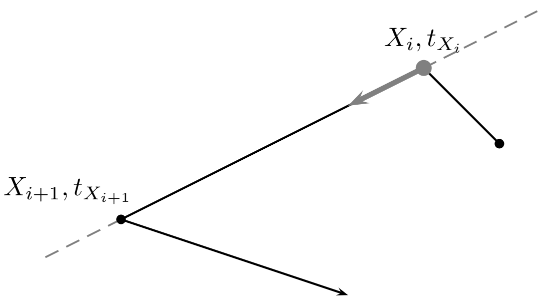


Graphic 2 – The path of an animal considered as a piecewise linear movement.

###### 2.3.3 Location of the rotating transects

The path of a clockwise rotating transect with time is defined by:

$$X_{t+\delta} = \begin{cases} X^1_{t+\delta} = -r\cos\left(\alpha_t + \dfrac{s\cdot\delta}{r}\right) + c \\ X^2_{t+\delta} = -r\sin\left(\alpha_t + \dfrac{s\cdot\delta}{r}\right) + c \end{cases} \tag{22}$$

with radius $r$, center $c$ and $\left(\alpha_t + \dfrac{s\cdot\delta}{r}\right) = 0$ at $t_0$.

###### 2.3.4 Error estimation

In order to compute $|\mathcal{S} - \widetilde{\mathcal{S}}_k|$, we aggregate the overall count in each of the two types of transects, meaning that we sum the counts for each lines of sight and speeds as

$$\sum\sum M_{\text{Linear transects}} = \sum\sum \begin{bmatrix} c_{11} & c_{12} & c_{13} & c_{14} \\ c_{21} & c_{22} & c_{23} & c_{24} \\ \dots & \dots & \dots & \dots \\ c_{61} & c_{62} & c_{63} & c_{64} \end{bmatrix}$$

with $c_{i,j}$ the aggregated count for line of sight $i$ and speed $j$. We then compute $\mathcal{S} - \widetilde{\mathcal{S}}_k$ as

$$|\mathcal{S} - \widetilde{\mathcal{S}}_k| = \sum\sum M_{\text{Linear transects (data)}} - \sum\sum M_{\text{Linear transects (simulation } k)}$$

We do the same for the rotating transects.

##### 2.4 Determinism

We investigate the determinism aspect of the statistics over a range of increasing $n_s$ values with $n_s = 10^4, 2\times 10^4, \dots, 4\times 10^5$. For each of those step values, a set of 100 BCR is simulated with parameters $p_I$, $p_F$ and $p_s$ estimated from the first deer (see Table 2). The following quantities are investigated:

1. the intra-bins variance in the normalized histograms of turning angles. For each $n_s$ value, the variance in each of the 20 bins is estimated and we use $\sum_1^{20} \mathrm{Var}(j)$ as an indicator of the overall intra-bins variance, with $\mathrm{Var}(j)$ being the variance of bin $j$.

2. the intra-density of the normalized home range area. Similarly to 1., we compute the variance of the estimated home range area for each density and for each $n_s$ value. Again, we sum the variances as an indicator of the overall variance as $n_s$ change. The home-range distributions are normalized such as the sum of their densities equals to 1.

3. the variance of the normalized distribution of the first 500 immobile transects counts.

4. the variance of the normalized count in mobile linear and in rotational transects. In order to assess the changes in counts as $n_s$ increase, we compute the variance of the overall counts for the different speeds and radiuses tested. Counts are normalized by the sum of the counts computed for all 100 simulations in every speed and radius.

5. the intra-disk size variance of the dilated paths. Similarly to 1. and for each $n_s$ value, we compute the variance of the estimated surface for each disk size. Again, we use proportions rather than absolute surfaces values: each surface is normalized by the sum of all the surfaces obtained after dilating with all different disk sizes. We sum the variances as an indicator of the overall variance as $n_s$ change.
