## Supplementary file 2 for "How to use random walks for modeling the movement of wild animals"

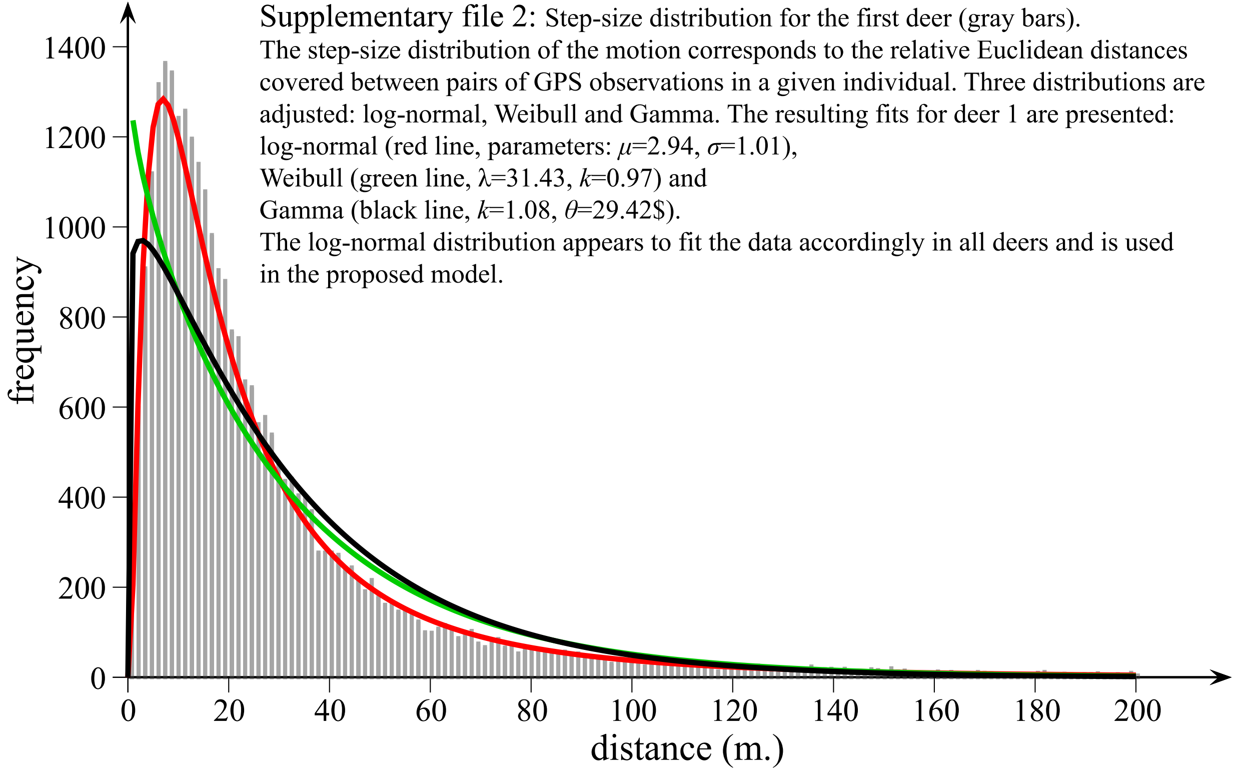


Supplementary file 2: Step-size distribution for the first deer (gray bars).
The step-size distribution of the motion corresponds to the relative Euclidean distances covered between pairs of GPS observations in a given individual. Three distributions are adjusted: log-normal, Weibull and Gamma. The resulting fits for deer 1 are presented:
log-normal (red line, parameters: $\mu$=2.94, $\sigma$=1.01),
Weibull (green line, $\lambda$=31.43, $k$=0.97) and
Gamma (black line, $k$=1.08, $\theta$=29.42$).
The log-normal distribution appears to fit the data accordingly in all deers and is used in the proposed model.
