## Supplementary file 3 for "How to use random walks for modeling the movement of wild animals"

Details of the 5 statistics. For each GPS data of deer 1 (left panels), a statistic is computed. Error estimates are detailed in the right panels.

### STATISTICS

### ERROR ESTIMATES

#### STAT 1 : Distribution of Turning Angles

Motion (data or simulated)

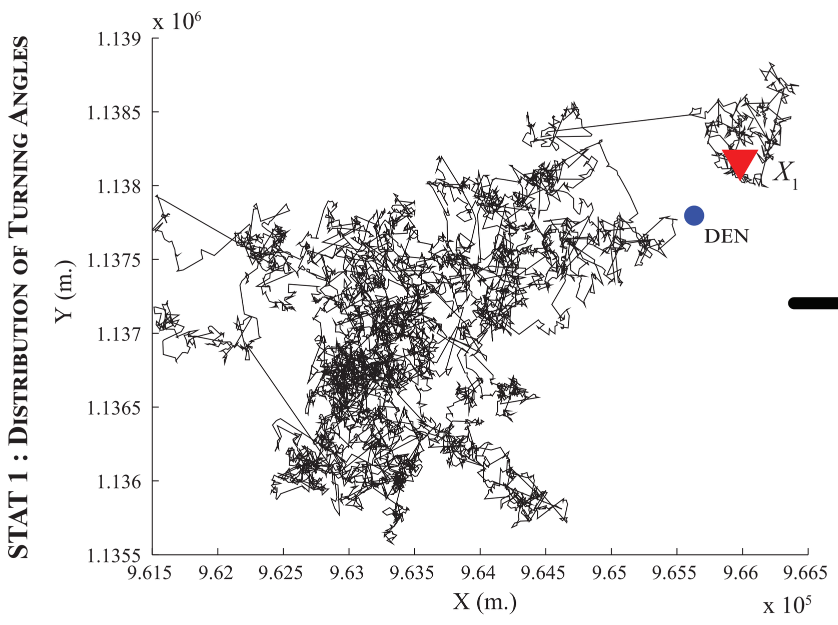


Angular distribution

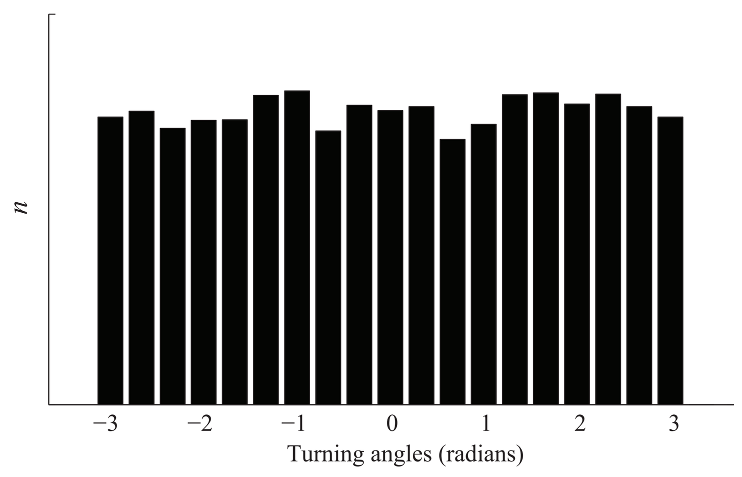


Difference between bins of the distribution

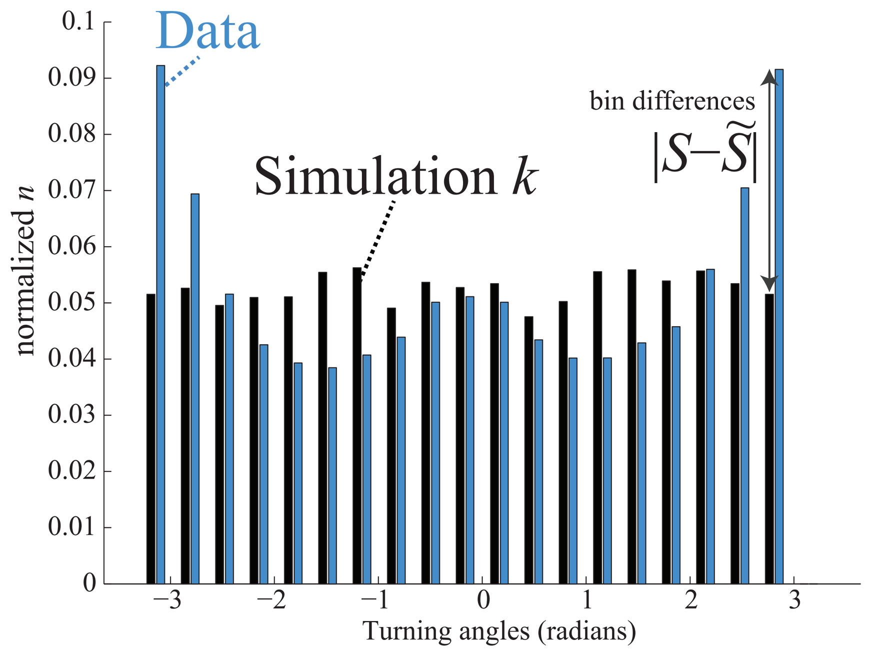


#### STAT 2 : Kernels Estimates

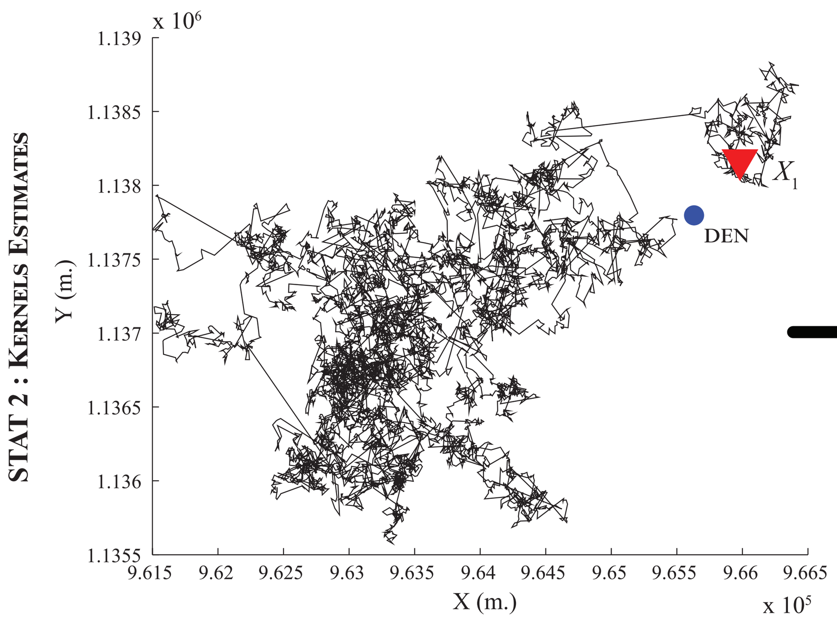


Home range (kernel estimate)
(over a mesh containing 1024 x 1024 nodes)

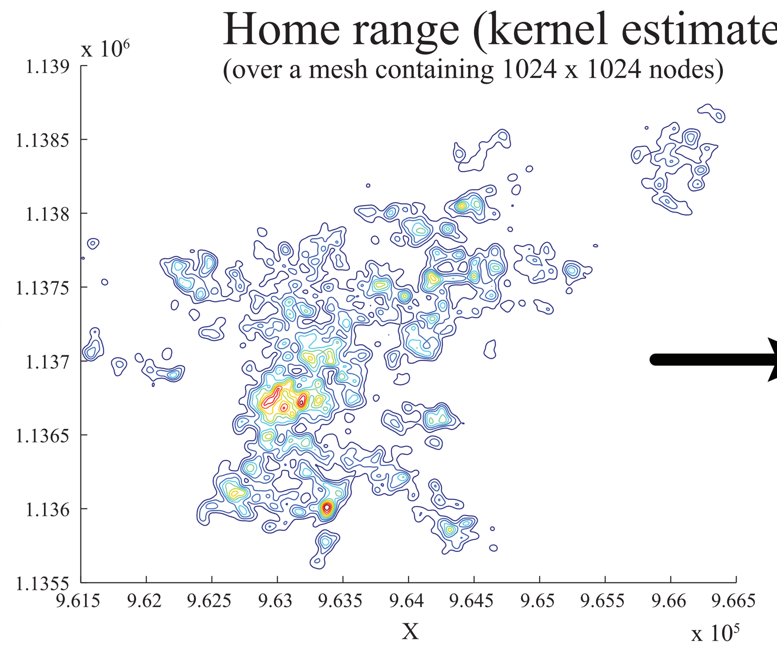


Collect the $n$ most dense nodes, representing $X$%* of the estimated kernel, then estimate the area covered by these $n$ nodes

*$X$ = [10, 20, 30, ..., 100]

Difference between estimated areas A for each $X$%:

| $X$ (%) | | 10 | 20 ... | 100 |
|---|---|---|---|---|
| $A_{data}$ | ($10^6$ m²) | 0.0074 | 0.0250 ... | 2.0331 |
| $A_{simulation\ k}$ | ($10^7$ m²) | 0.0127 | 0.0310 ... | 3.6114 |
| $\|S-\tilde{S}\|$ | ($10^7$ m²) | 0.0120 | 0.0285 ... | 3.4081 |

#### STAT 3 : Dilation

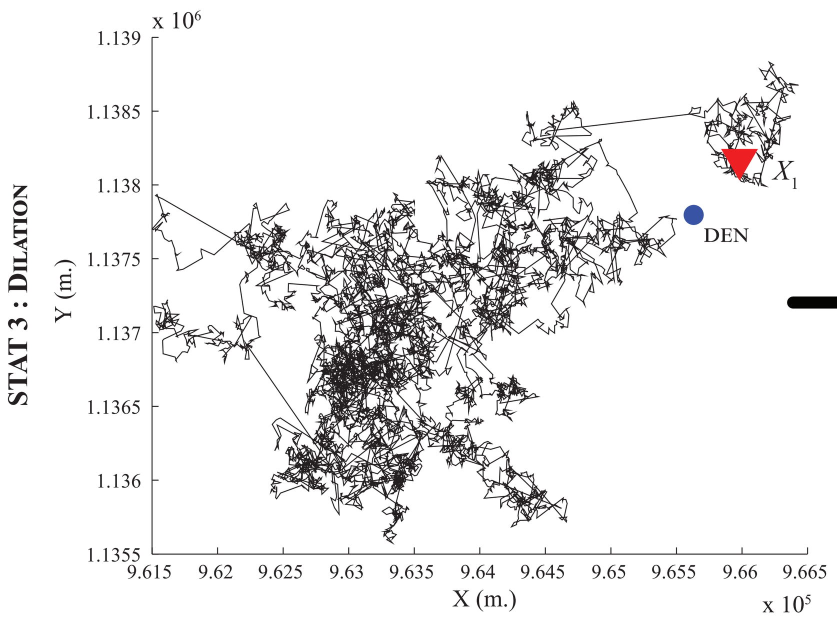


Dilation
(using a disk of size 1, 2, ..., 100)

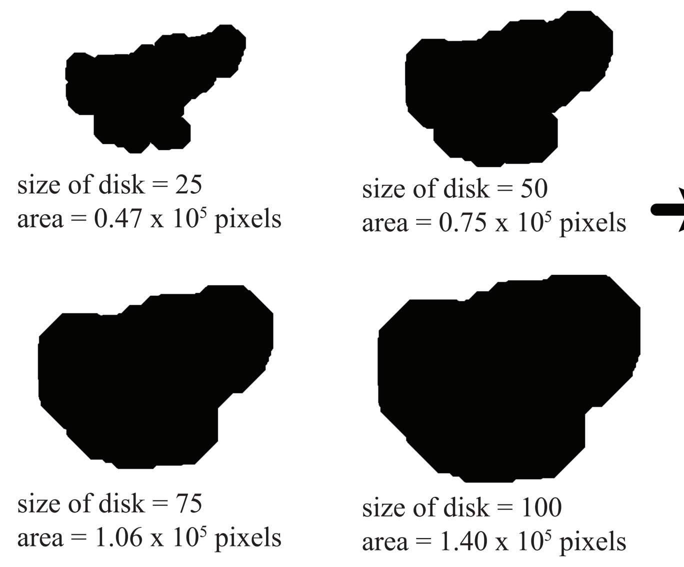


Collect the estimated areas for all sizes of the disk

Difference between estimated areas A:

| size of disk (pixels) | | ... | 25 | ... | 50 | ... | 100 |
|---|---|---|---|---|---|---|---|
| $A_{data}$ | ($10^5$ pixels) | ... | 0.4080 | ... | 0.6577 | ... | 1.2655 |
| $A_{simulation\ k}$ | ($10^5$ pixels) | ... | 0.4717 | ... | 0.7488 | ... | 1.4027 |
| $\|S-\tilde{S}\|$ | ($10^4$ pixels) | ... | 0.6366 | ... | 0.9103 | ... | 1.3716 |

#### STAT 4 : Immobile Transects

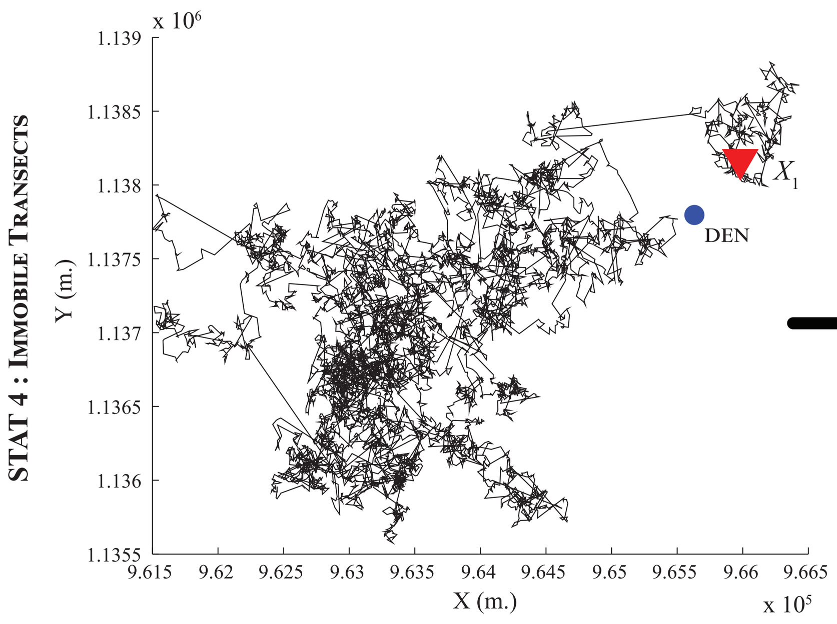


Immobile transects
(with fixed area of sight)

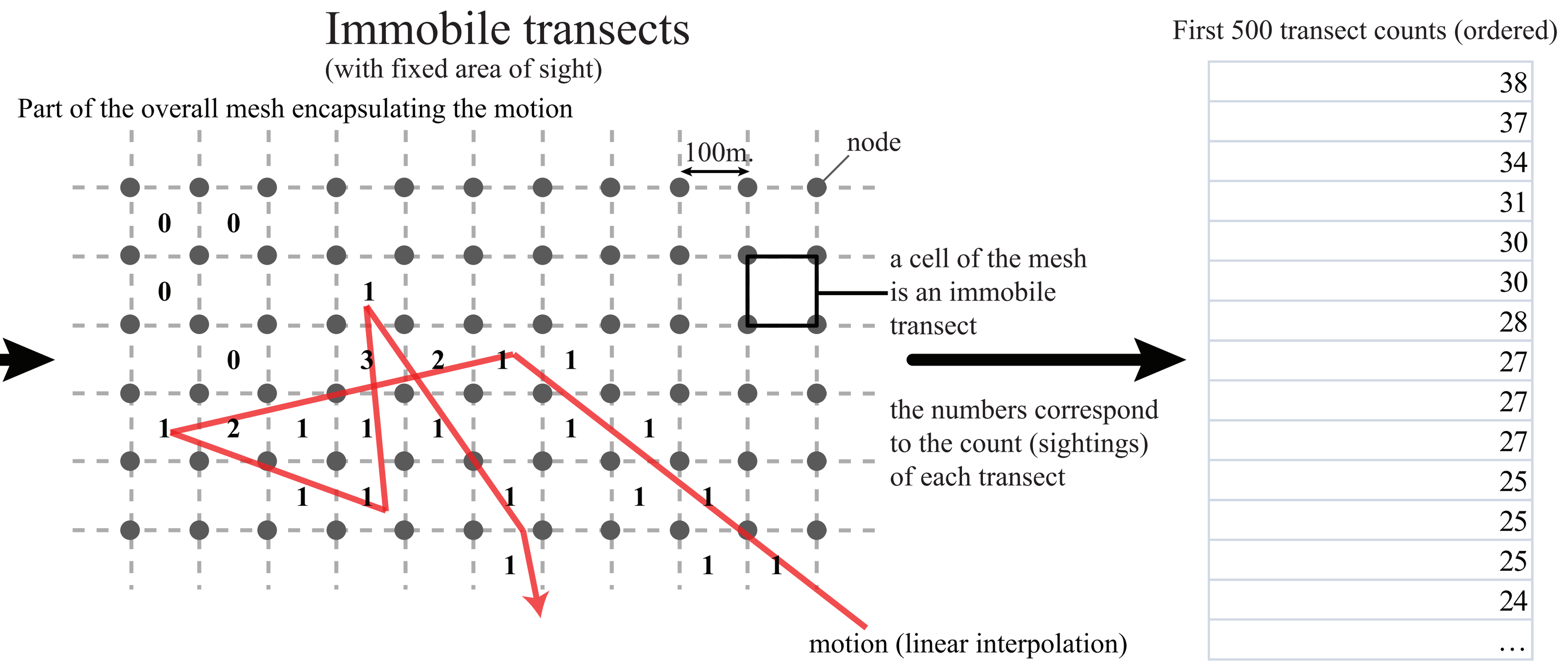


Difference between first 500 transect counts (ordered)

| data | simulation $k$ | $\|S-\tilde{S}\|$ |
|---|---|---|
| 388 | 38 | 350 |
| 380 | 37 | 343 |
| 309 | 34 | 275 |
| 258 | 31 | 227 |
| 227 | 30 | 197 |
| 225 | 30 | 195 |
| ... | ... | ... |

#### STAT 5 : Mobile Transects (Linear)

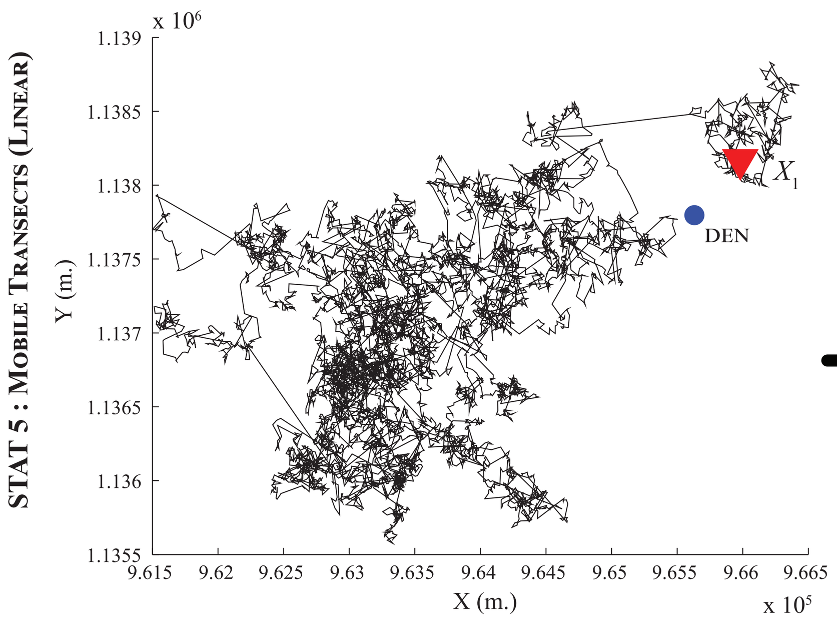


Mobile transects (linear)
(fixed area of sight, constant speed)
δ = (average speed of the animal)/2

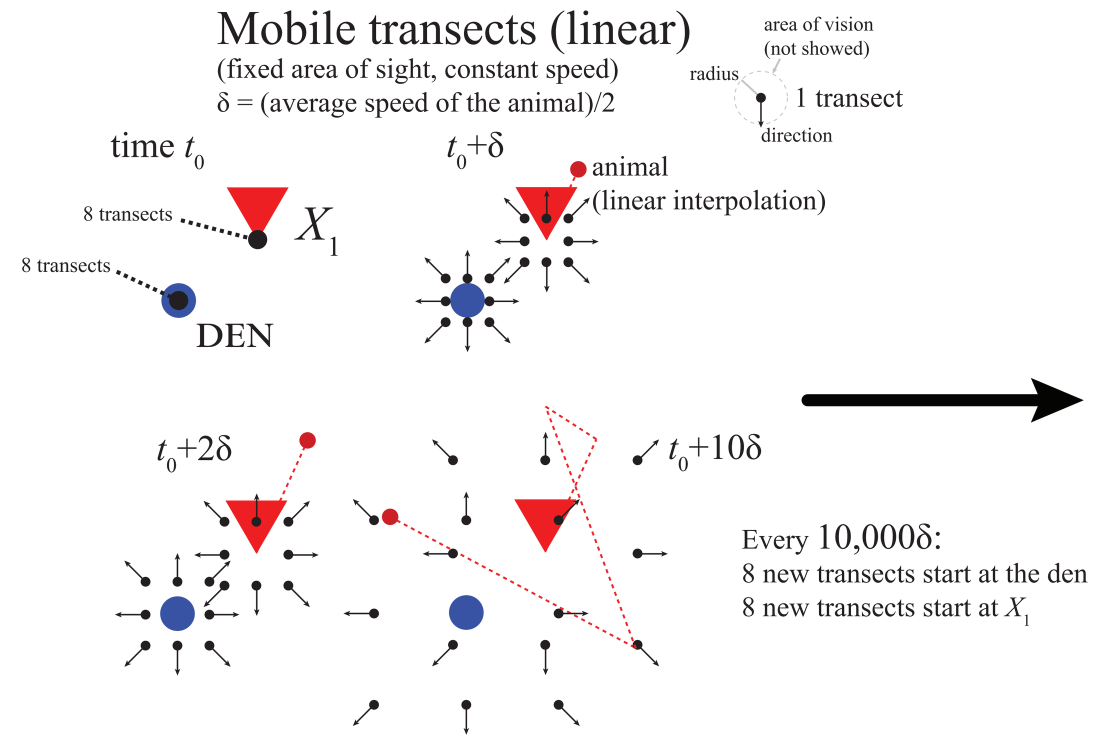


For each speed and radius, all transects counts are aggregated

Difference between transect counts for all radius and speeds:

$|S-\tilde{S}|$ = |Total count (data) - total count (simulation $k$)|

#### STAT 5 : Mobile Transects (Rotating)

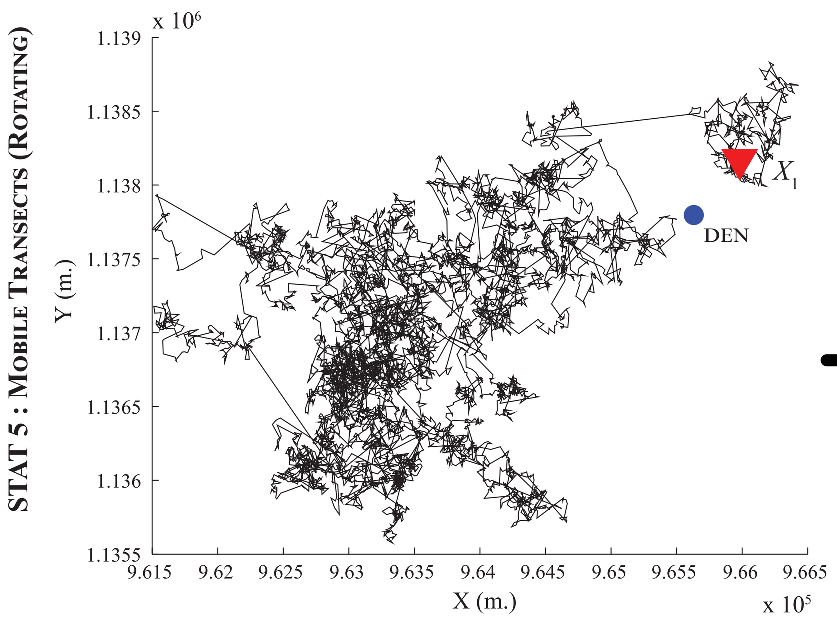


Mobile transects (rotating)
(fixed area of sight, constant speed)
δ = (average speed of the animal)/2

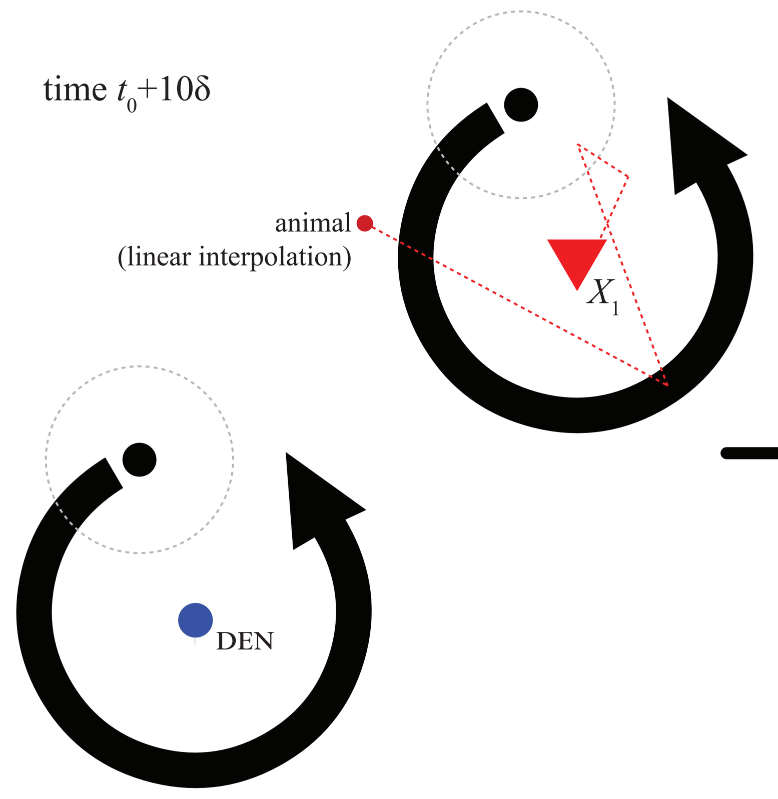


For each speed and radius, the two transects counts are aggregated

Difference between transect counts for all radius and speeds:

$|S-\tilde{S}|$ = |Total count (data) - total count (simulation $k$)|
