## Supplementary file 4 for "How to use random walks for modeling the movement of wild animals"

Density of error $e$ of all 5 configurations tested in each statistic for all deers. Densities are fitted by the Epanechnikov kernel function.

### Distribution of turning angles

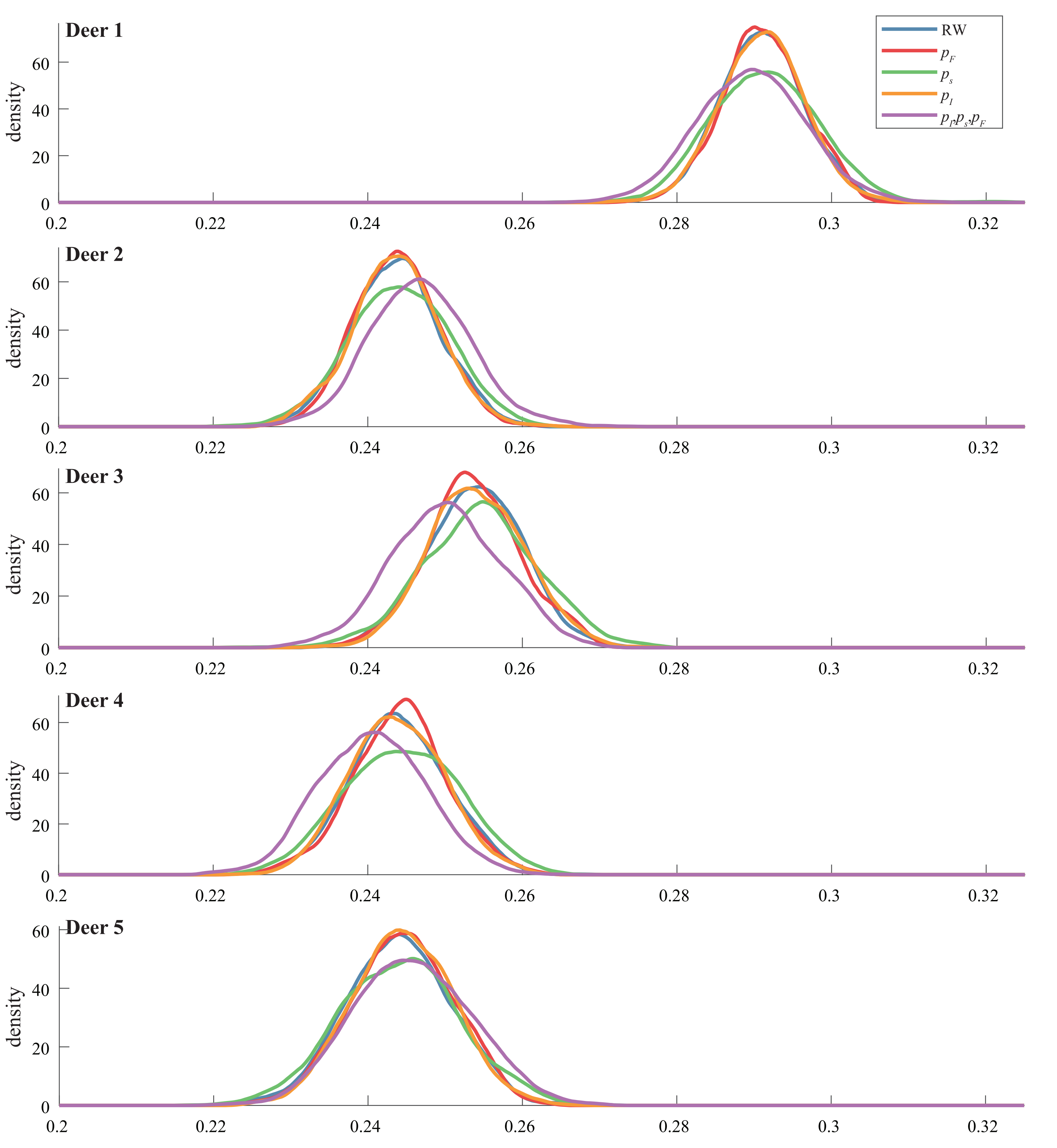


### Home range

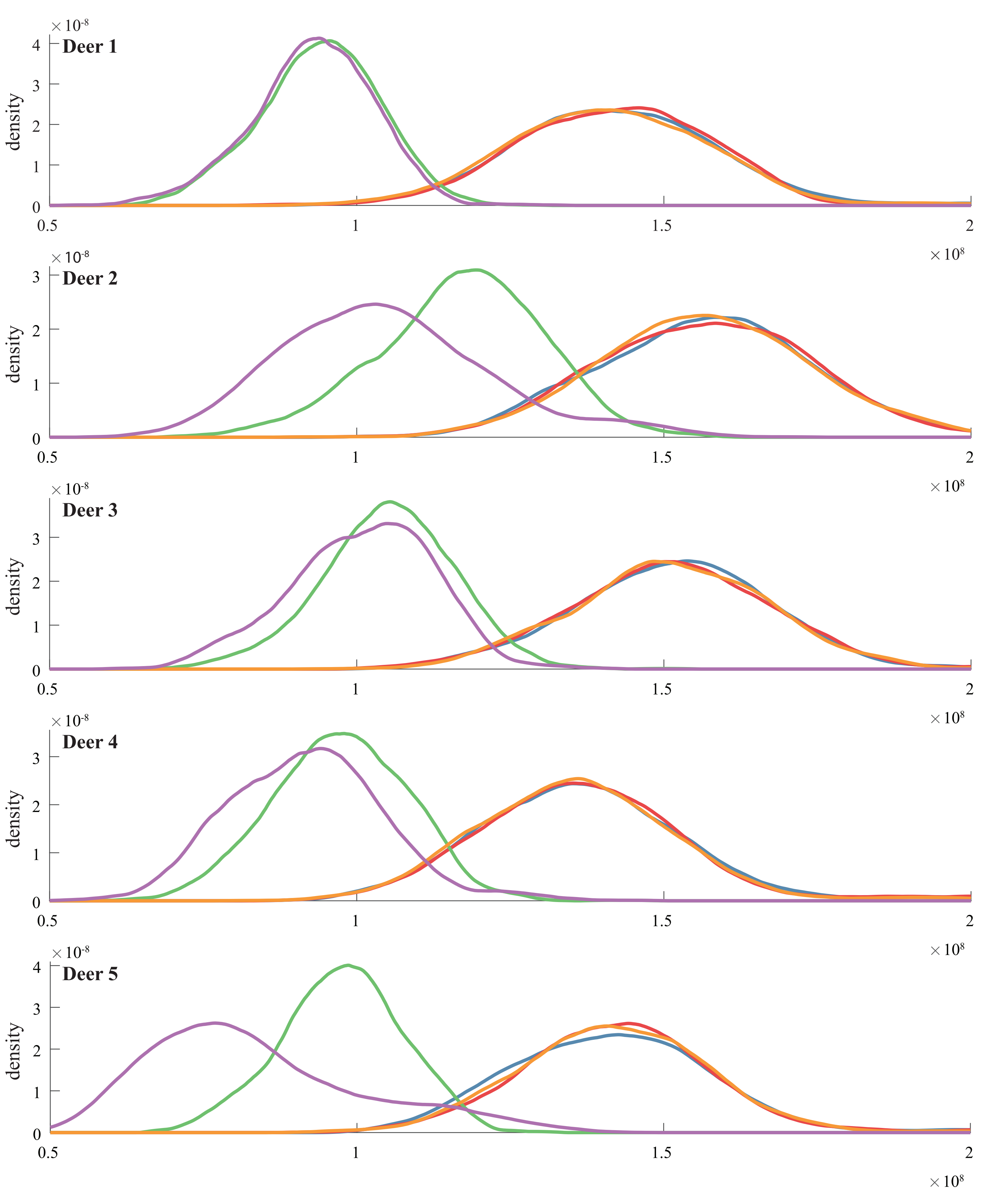


### Dilation

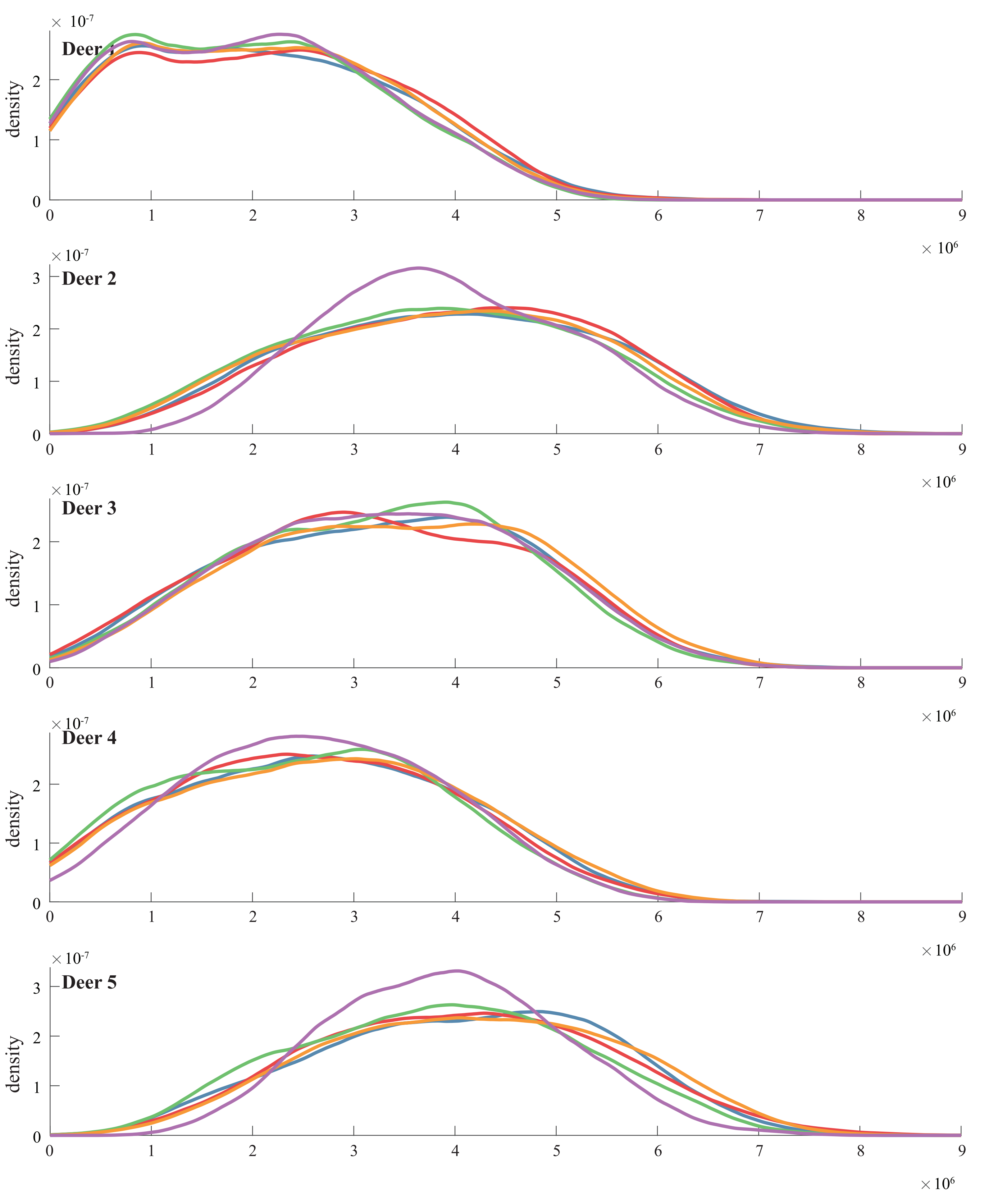


### Immobile transects

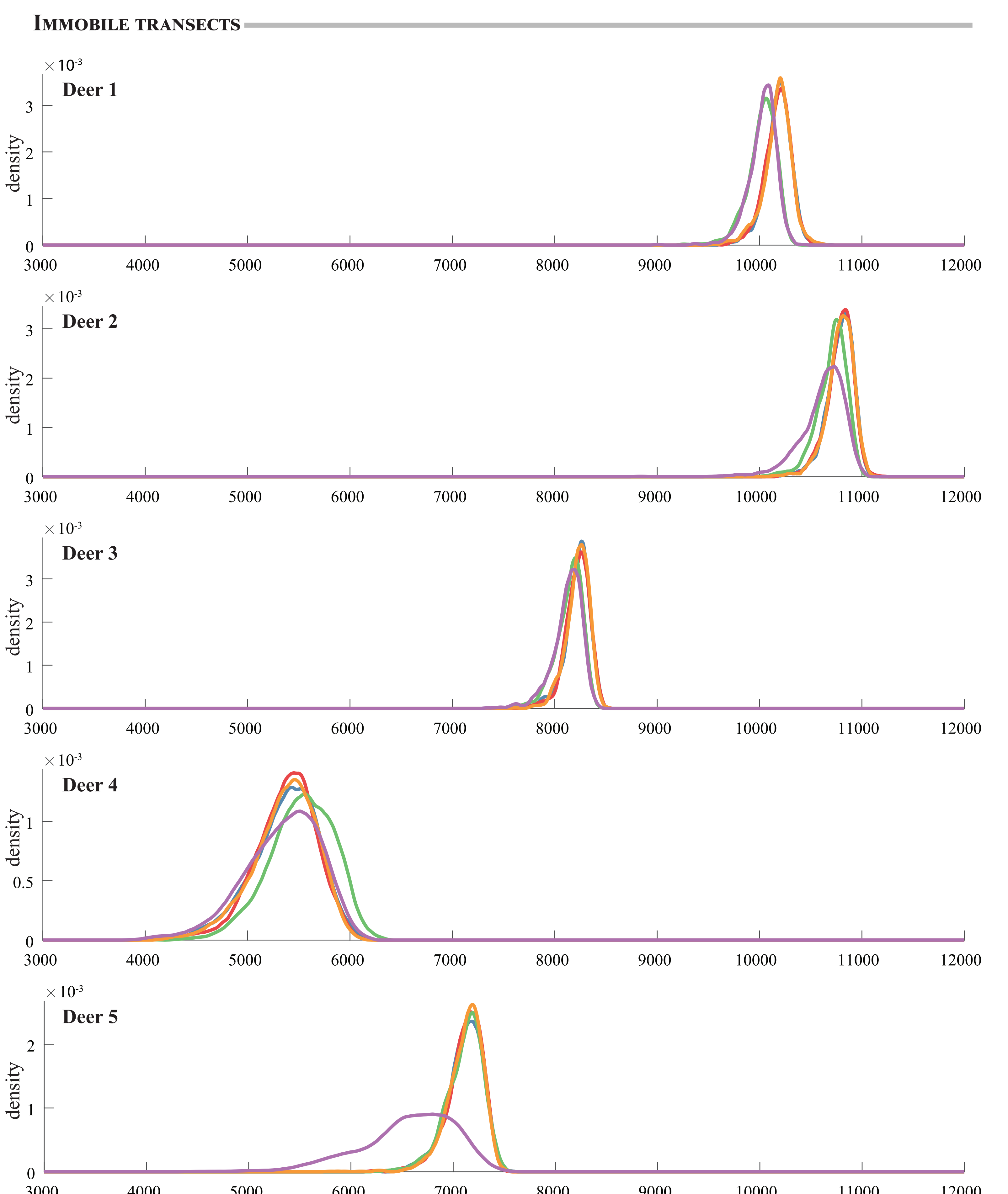


### Mobile transects (linear)

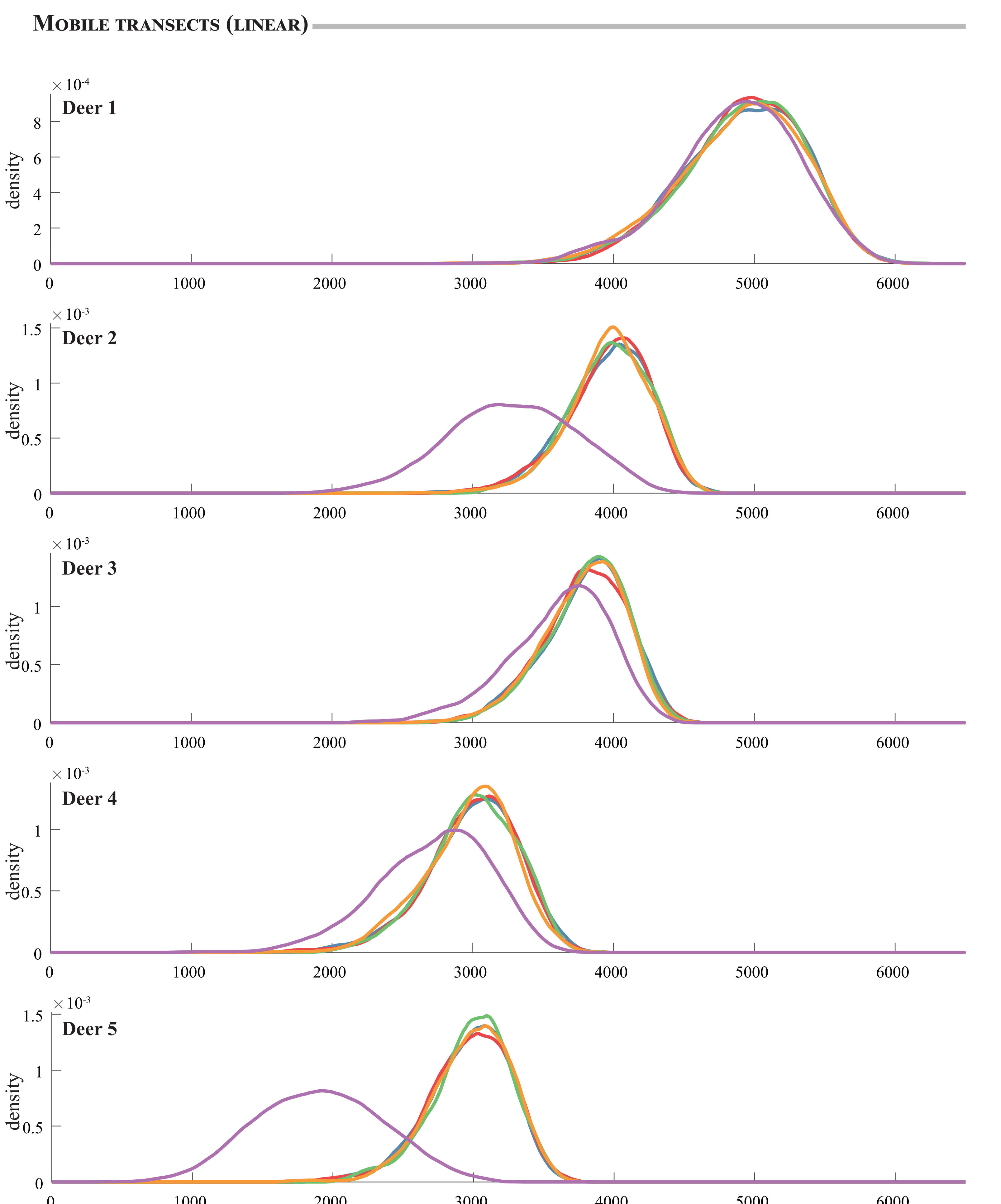


### Mobile transects (rotary)

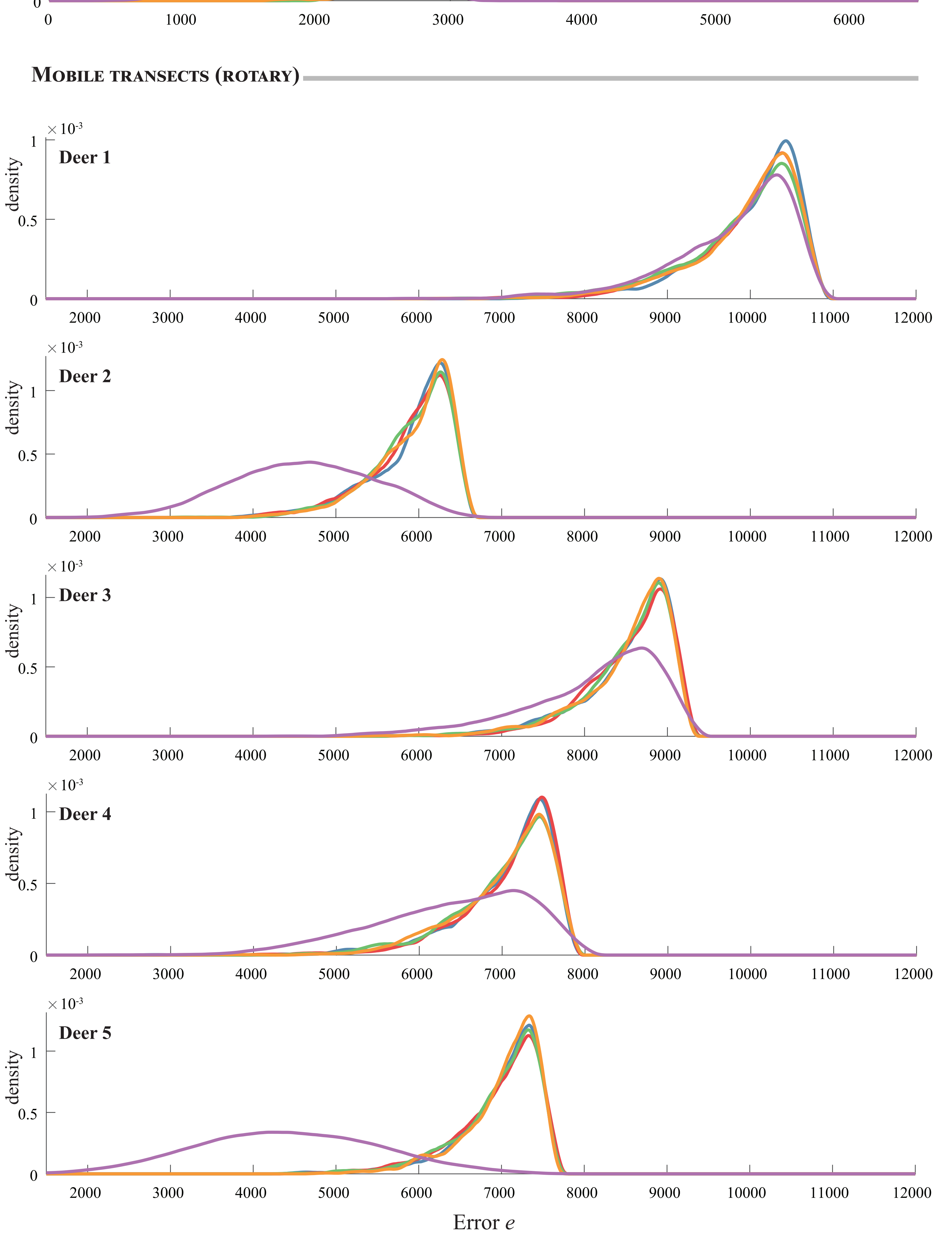
