## Supplementary file 5 for "How to use random walks for modeling the movement of wild animals"

Variance of the statistics with an increasing number of simulated steps $n_s$, from $n_s=10^4$ to $n_s=4\times10^5$. We perform additional simulations in the $[10^4, 2\times10^4]$ interval to show the rapid decrease in the home range statistic (**B**). We also show the intra-class variance for all 100 disks sizes in the dilation (**C**) and the intra-bins variance for each of the 500 bins of each distribution in immobile transects (**D**).
The variance of the normalized count in mobile transects are detailed for all 4 speeds and 8 radiuses tested for mobile transects (**E** and **F**).

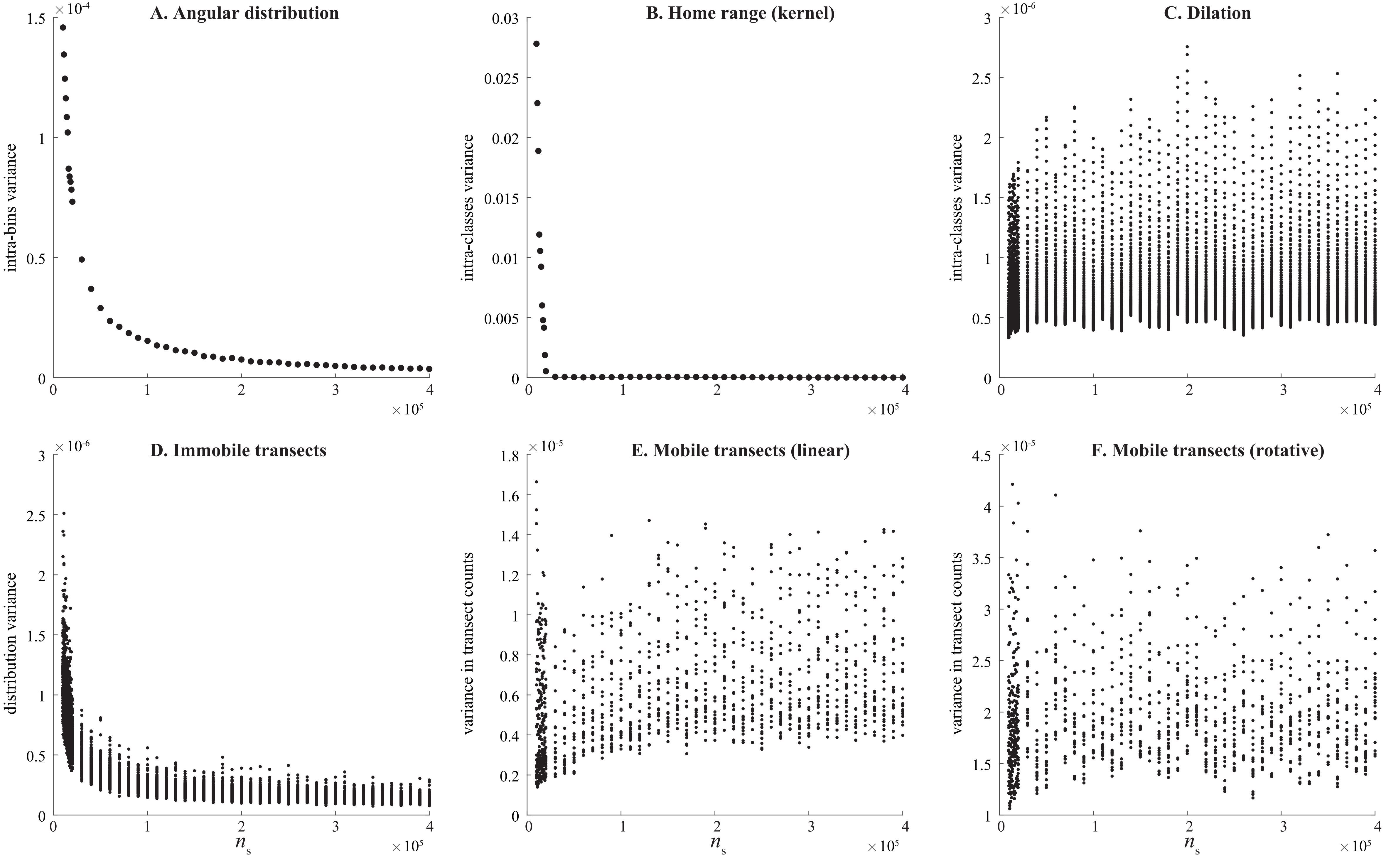
