## Supplementary file 6 for "How to use random walks for modeling the movement of wild animals"

The values of $p_I$, $p_F$ and $p_s$ are estimated with increasing subsampling (or decimation) rate $k$ for each deer: panels **A** - **E**. The first X-axis corresponds to the subsampling rate $k$ while the second X-axis is $\overline{T}$, the average sampling time. An example of subsampling is presented in panel **F**, with increasing subsampling rate $k$ from $k$=1 (upper left sub panel) to $k$=10 (lower right sub panel).

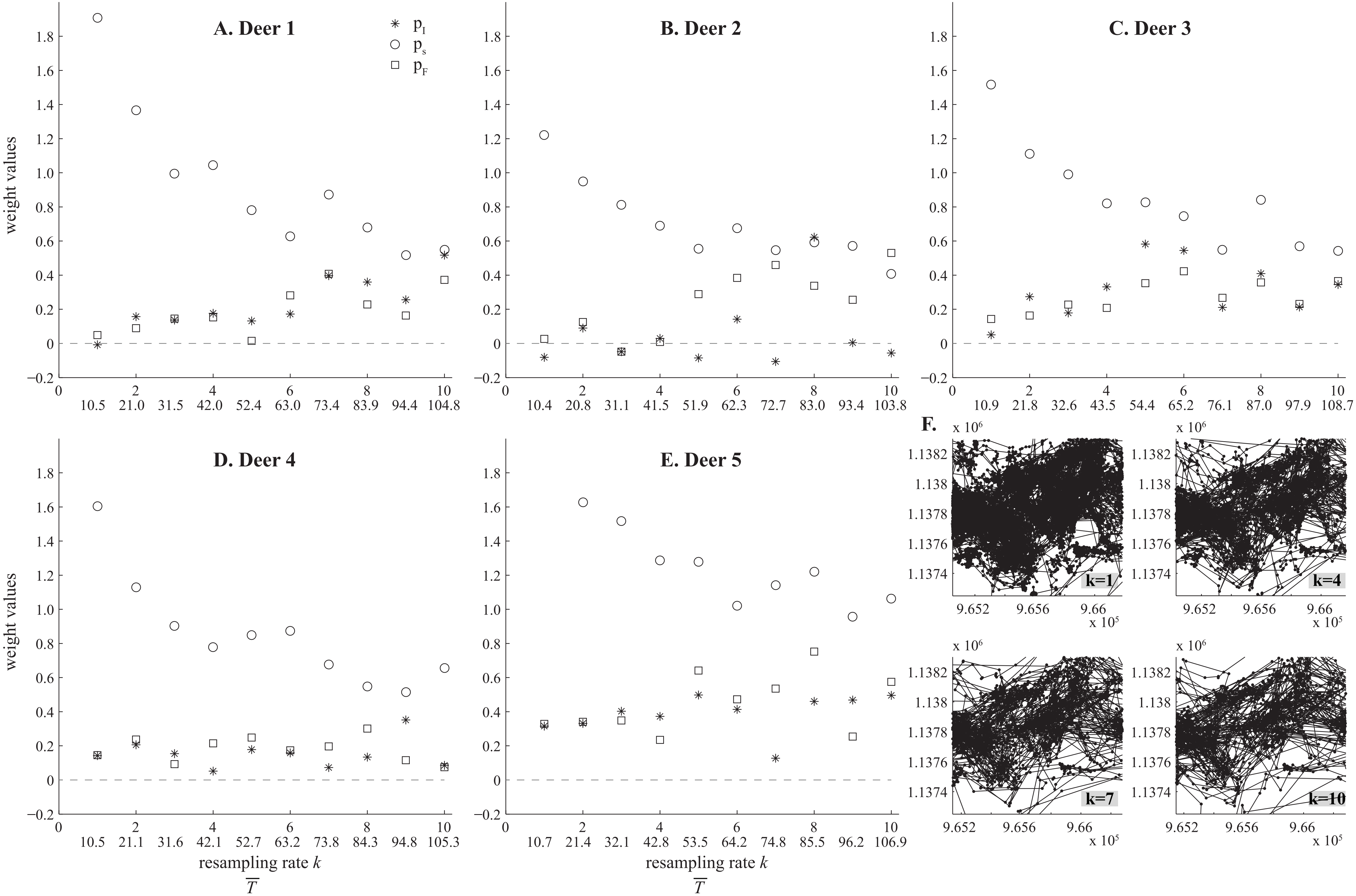
