## Supplementary file 7 for "How to use random walks for modeling the movement of wild animals"

The alpha shape using a fixed alpha radius of 60m are presented for all deers (green areas). Only the localizations near the isobarycenter are displayed here and distant localizations are not showed. Voids of interest are near the center of the shapes, and labelled with circled numbers.

Deer 1

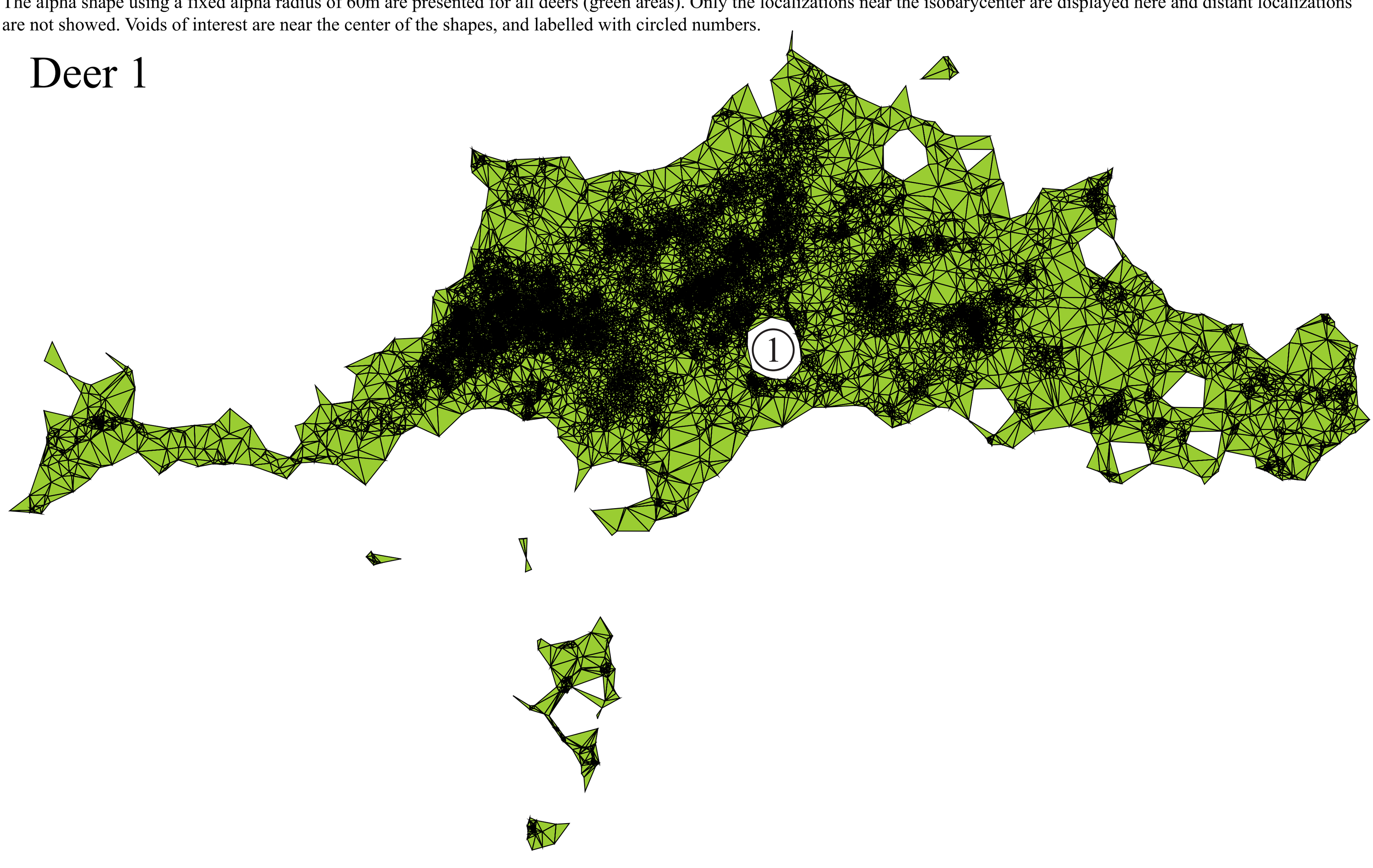


Deer 2

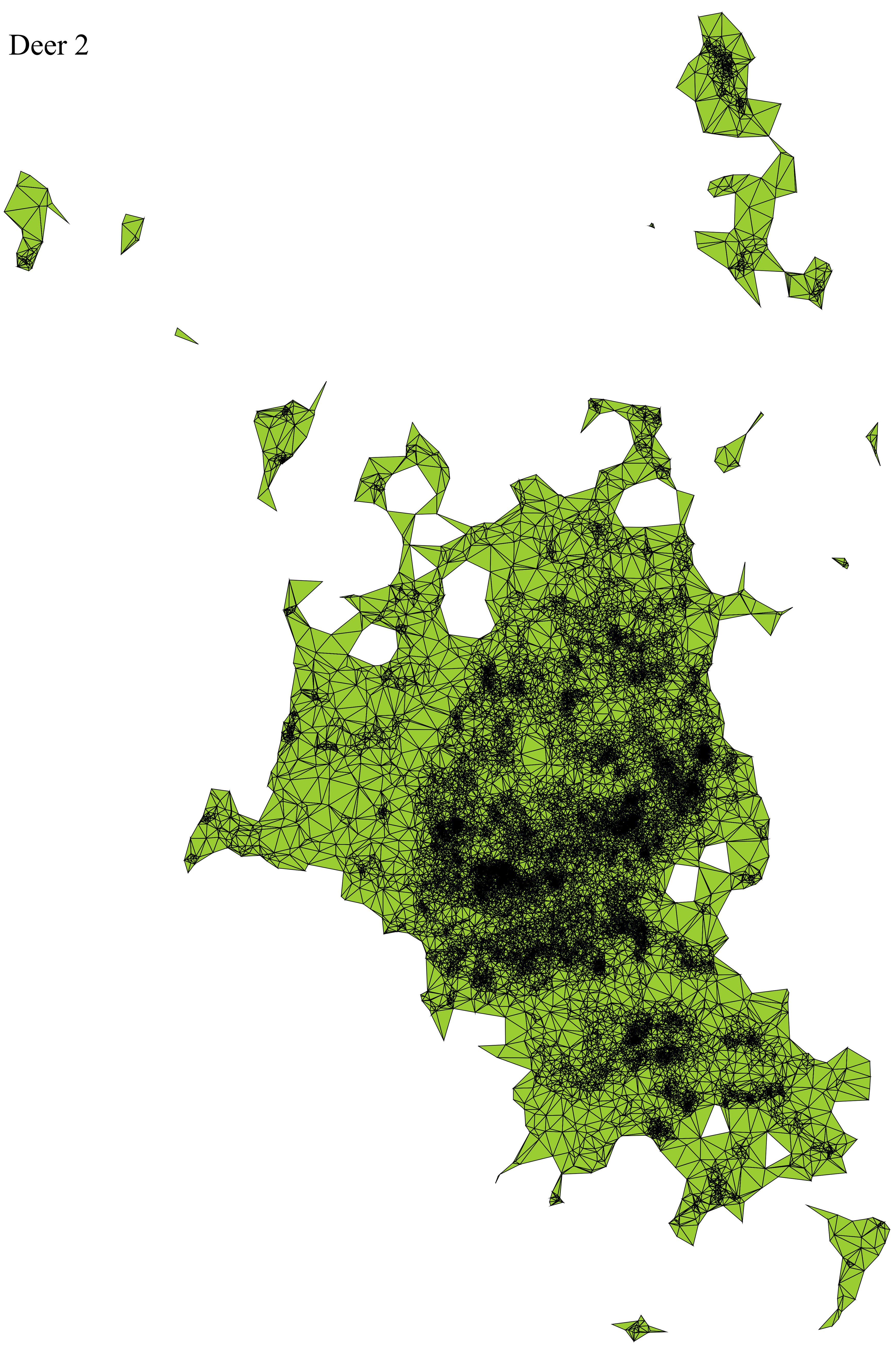

Deer 3

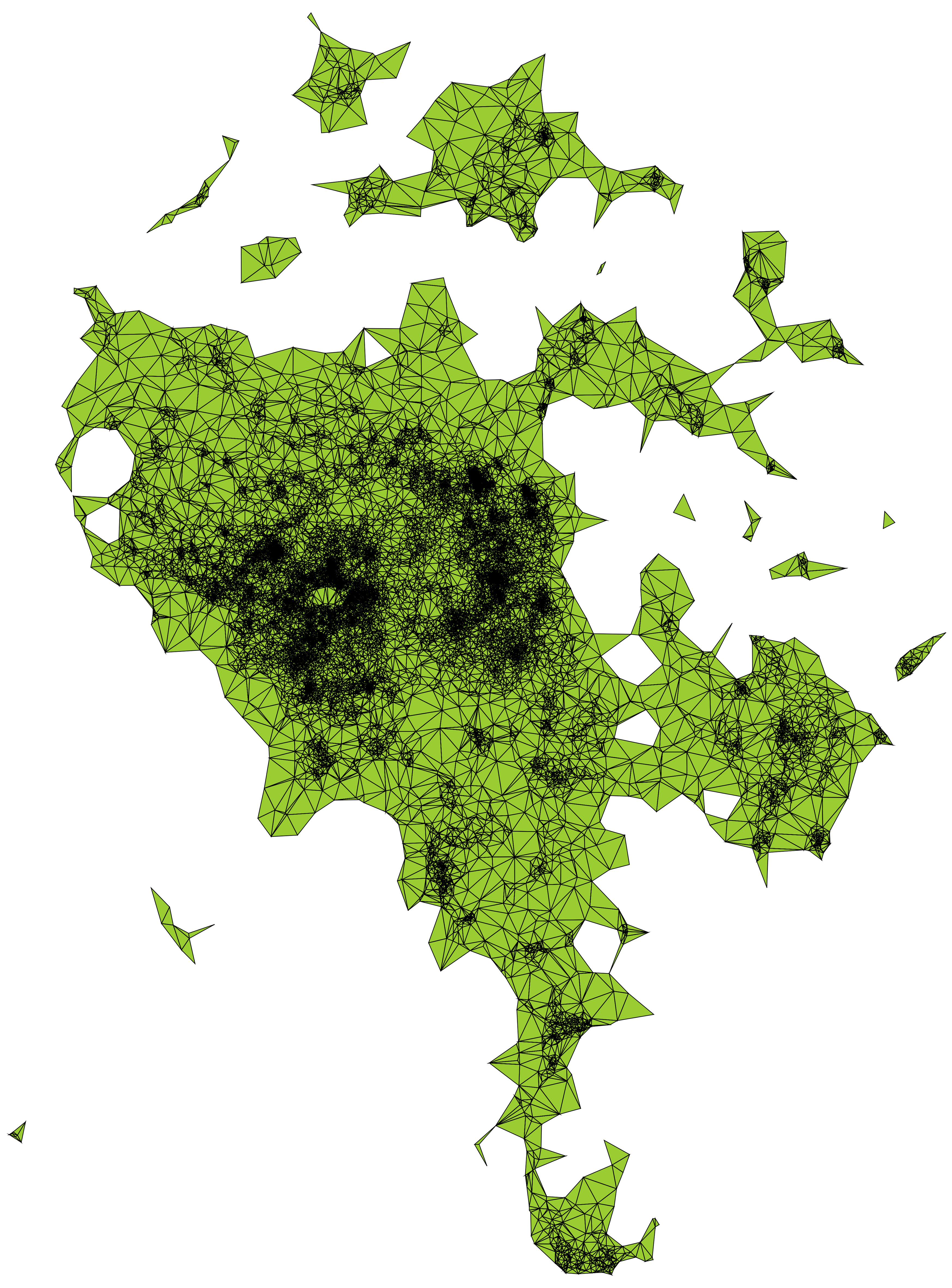

Deer 4

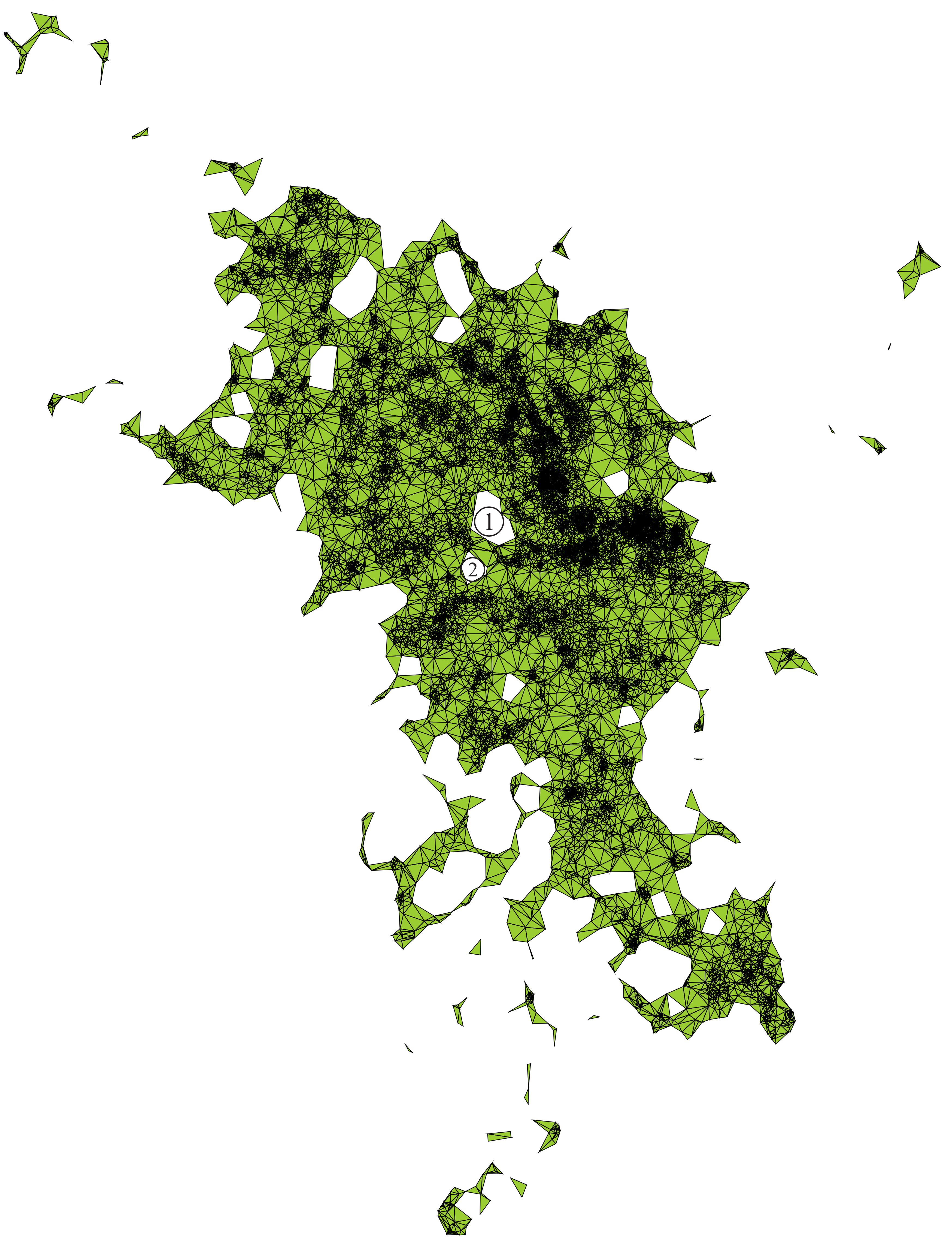


Deer 5

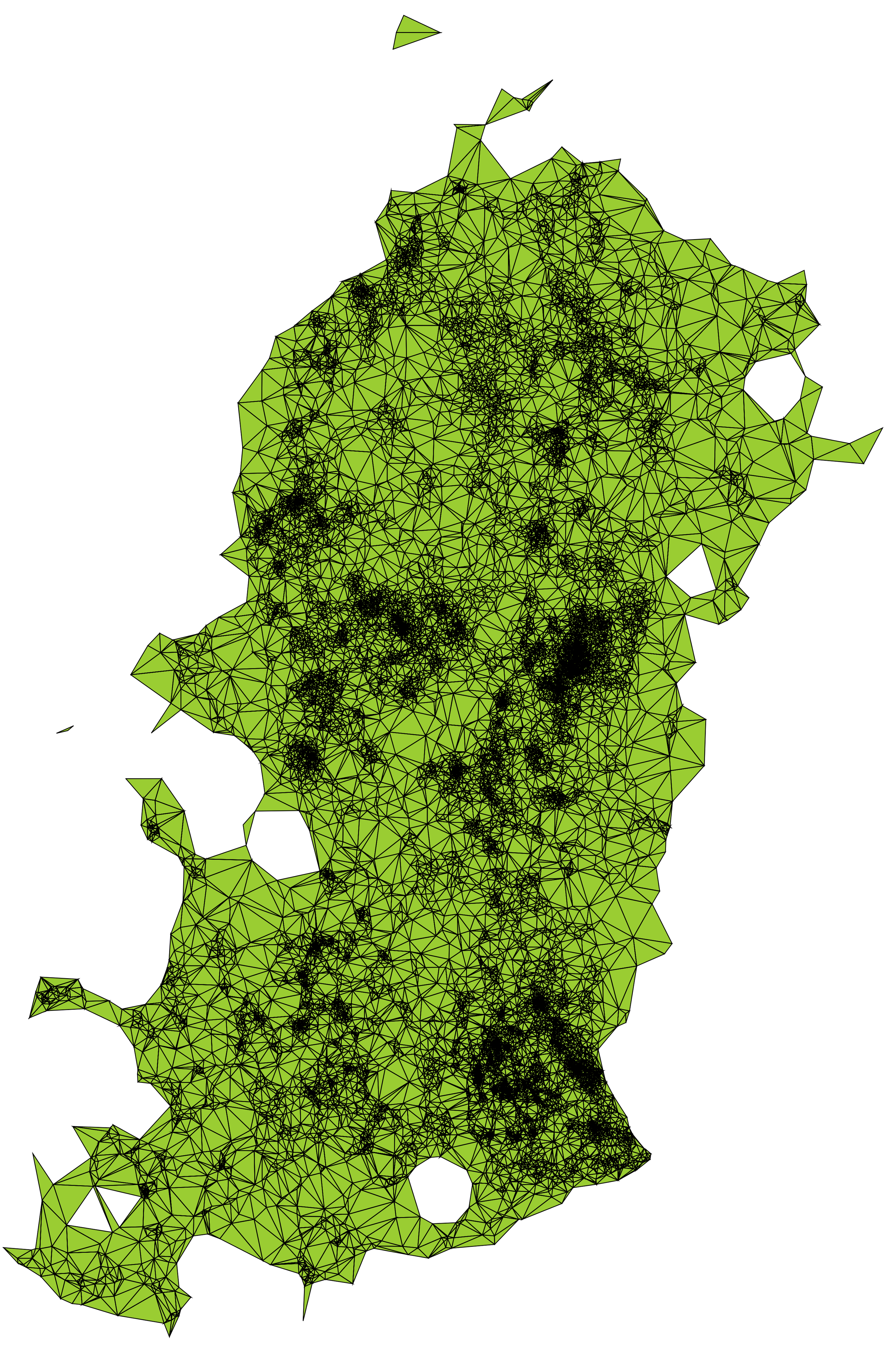
