## Supplementary file 8 - Table S1 for "How to use random walks for modeling the movement of wild animals"

Parameters values when using only one parameter instead of the three.

| **Parameters configuration** | **Deer** | $p_I$ | $p_s$ | $p_F$ |
|---|---|---|---|---|
| $p_I$ | Deer#1 | -0.1742 | | |
| $p_I$ | Deer#2 | -0.1176 | | |
| $p_I$ | Deer#3 | -0.0928 | | |
| $p_I$ | Deer#4 | -0.0774 | | |
| $p_I$ | Deer#5 | -0.0402 | | |
| $p_s$ | Deer#1 | | 1.9718 | |
| $p_s$ | Deer#2 | | 1.3956 | |
| $p_s$ | Deer#3 | | 1.6534 | |
| $p_s$ | Deer#4 | | 1.4871 | |
| $p_s$ | Deer#5 | | 1.6207 | |
| $p_F$ | Deer#1 | | | -0.1746 |
| $p_F$ | Deer#2 | | | -0.0694 |
| $p_F$ | Deer#3 | | | -0.1474 |
| $p_F$ | Deer#4 | | | -0.11 |
| $p_F$ | Deer#5 | | | -0.0164 |
| $p_I$, $p_s$, $p_F$ | Deer#1 | 0.0111 | 2.0073 | 0.0121 |
| $p_I$, $p_s$, $p_F$ | Deer#2 | 0.0643 | 1.4367 | 0.1316 |
| $p_I$, $p_s$, $p_F$ | Deer#3 | 0.1174 | 1.697 | 0.0531 |
| $p_I$, $p_s$, $p_F$ | Deer#4 | 0.101 | 1.5157 | 0.063 |
| $p_I$, $p_s$, $p_F$ | Deer#5 | 0.2176 | 1.6606 | 0.2382 |
