## Supplementary file 9 - Table S2 for "How to use random walks for modeling the movement of wild animals"

Results of the statistics showing model performance when using only one parameter instead of the three.

| Deer# | Stat | Configuration | mean($e$) | std($e$) | median($e$) | interquartile($e$) |
|---|---|---|---|---|---|---|
| 1 | Distribution of turning angles | RW | 0.2910 | 0.0053 | 0.2909 | 0.0072 |
| 1 | Distribution of turning angles | $p_F$ | 0.2912 | 0.0052 | 0.2910 | 0.0070 |
| 1 | Distribution of turning angles | $p_s$ | 0.2914 | 0.0068 | 0.2912 | 0.0093 |
| 1 | Distribution of turning angles | $p_I$ | 0.2910 | 0.0053 | 0.2912 | 0.0071 |
| 1 | Distribution of turning angles | $p_I, p_s, p_F$ | 0.2896 | 0.0068 | 0.2898 | 0.0093 |
| 1 | Home range (Kernel estimate) | RW | 1.42E+08 | 1.67E+07 | 1.42E+08 | 2.15E+07 |
| 1 | Home range (Kernel estimate) | $p_F$ | 1.43E+08 | 1.65E+07 | 1.43E+08 | 2.14E+07 |
| 1 | Home range (Kernel estimate) | $p_s$ | 9.40E+07 | 9.65E+06 | 9.44E+07 | 1.30E+07 |
| 1 | Home range (Kernel estimate) | $p_I$ | 1.42E+08 | 1.74E+07 | 1.41E+08 | 2.20E+07 |
| 1 | Home range (Kernel estimate) | $p_I, p_s, p_F$ | 9.29E+07 | 9.85E+06 | 9.34E+07 | 1.24E+07 |
| 1 | Dilation | RW | 2.08E+06 | 1.28E+06 | 2.00E+06 | 2.08E+06 |
| 1 | Dilation | $p_F$ | 2.14E+06 | 1.28E+06 | 2.14E+06 | 2.11E+06 |
| 1 | Dilation | $p_s$ | 1.97E+06 | 1.21E+06 | 1.94E+06 | 1.93E+06 |
| 1 | Dilation | $p_I$ | 2.09E+06 | 1.24E+06 | 2.08E+06 | 1.97E+06 |
| 1 | Dilation | $p_I, p_s, p_F$ | 2.01E+06 | 1.21E+06 | 1.96E+06 | 1.92E+06 |
| 1 | Immobile transects | RW | 10178.0 | 132.8 | 10189.0 | 155.0 |
| 1 | Immobile transects | $p_F$ | 10174.0 | 130.1 | 10190.0 | 165.5 |
| 1 | Immobile transects | $p_s$ | 10023.0 | 142.6 | 10041.0 | 171.0 |
| 1 | Immobile transects | $p_I$ | 10175.0 | 140.2 | 10190.0 | 152.5 |
| 1 | Immobile transects | $p_I, p_s, p_F$ | 10029.0 | 137.3 | 10051.0 | 160.5 |
| 1 | Mobile transects (linear) | RW | 4916.7 | 424.0 | 4956.5 | 605.5 |
| 1 | Mobile transects (linear) | $p_F$ | 4923.6 | 412.8 | 4957.5 | 555.5 |
| 1 | Mobile transects (linear) | $p_s$ | 4922.4 | 423.8 | 4961.5 | 561.0 |
| 1 | Mobile transects (linear) | $p_I$ | 4898.4 | 438.1 | 4948.5 | 601.0 |
| 1 | Mobile transects (linear) | $p_I, p_s, p_F$ | 4876.2 | 432.4 | 4909.0 | 567.0 |
| 1 | Mobile transects (rotating) | RW | 9994.3 | 594.0 | 10169.0 | 737.5 |
| 1 | Mobile transects (rotating) | $p_F$ | 9948.4 | 632.3 | 10138.0 | 757.0 |
| 1 | Mobile transects (rotating) | $p_s$ | 9876.4 | 690.7 | 10063.0 | 833.5 |
| 1 | Mobile transects (rotating) | $p_I$ | 9946.3 | 642.3 | 10128.0 | 735.0 |
| 1 | Mobile transects (rotating) | $p_I, p_s, p_F$ | 9796.4 | 727.2 | 10009.0 | 939.5 |
| 2 | Distribution of turning angles | RW | 0.2436 | 0.0057 | 0.2437 | 0.0074 |
| 2 | Distribution of turning angles | $p_F$ | 0.2437 | 0.0053 | 0.2436 | 0.0074 |
| 2 | Distribution of turning angles | $p_s$ | 0.2441 | 0.0064 | 0.2441 | 0.0091 |
| 2 | Distribution of turning angles | $p_I$ | 0.2436 | 0.0056 | 0.2437 | 0.0072 |
| 2 | Distribution of turning angles | $p_I, p_s, p_F$ | 0.2468 | 0.0065 | 0.2467 | 0.0089 |
| 2 | Home range (Kernel estimate) | RW | 1.58E+08 | 2.02E+07 | 1.58E+08 | 2.37E+07 |
| 2 | Home range (Kernel estimate) | $p_F$ | 1.58E+08 | 1.99E+07 | 1.57E+08 | 2.44E+07 |
| 2 | Home range (Kernel estimate) | $p_s$ | 1.17E+08 | 1.33E+07 | 1.18E+08 | 1.64E+07 |
| 2 | Home range (Kernel estimate) | $p_I$ | 1.58E+08 | 1.90E+07 | 1.57E+08 | 2.34E+07 |
| 2 | Home range (Kernel estimate) | $p_I, p_s, p_F$ | 1.05E+08 | 1.68E+07 | 1.04E+08 | 2.19E+07 |
| 2 | Dilation | RW | 4.00E+06 | 1.45E+06 | 4.00E+06 | 2.32E+06 |
| 2 | Dilation | $p_F$ | 4.02E+06 | 1.40E+06 | 4.10E+06 | 2.13E+06 |
| 2 | Dilation | $p_s$ | 3.80E+06 | 1.42E+06 | 3.84E+06 | 2.20E+06 |
| 2 | Dilation | $p_I$ | 3.91E+06 | 1.44E+06 | 3.94E+06 | 2.26E+06 |
| 2 | Dilation | $p_I, p_s, p_F$ | 3.92E+06 | 1.18E+06 | 3.83E+06 | 1.77E+06 |
| 2 | Immobile transects | RW | 10783.0 | 135.7 | 10804.0 | 159.0 |
| 2 | Immobile transects | $p_F$ | 10784.0 | 134.6 | 10804.0 | 158.5 |
| 2 | Immobile transects | $p_s$ | 10705.0 | 148.7 | 10733.0 | 186.0 |
| 2 | Immobile transects | $p_I$ | 10786.0 | 131.4 | 10800.0 | 161.0 |
| 2 | Immobile transects | $p_I, p_s, p_F$ | 10617.0 | 219.8 | 10658.0 | 259.0 |
| 2 | Mobile transects (linear) | RW | 3955.0 | 297.3 | 3986.5 | 398.5 |
| 2 | Mobile transects (linear) | $p_F$ | 3961.4 | 294.0 | 3997.0 | 379.5 |
| 2 | Mobile transects (linear) | $p_s$ | 3973.9 | 280.9 | 3994.0 | 387.5 |
| 2 | Mobile transects (linear) | $p_I$ | 3969.3 | 282.9 | 3988.5 | 367.0 |
| 2 | Mobile transects (linear) | $p_I, p_s, p_F$ | 3277.0 | 445.7 | 3287.5 | 627.5 |
| 2 | Mobile transects (rotating) | RW | 5911.9 | 495.9 | 6059.5 | 610.5 |
| 2 | Mobile transects (rotating) | $p_F$ | 5899.6 | 481.5 | 6020.5 | 608.5 |
| 2 | Mobile transects (rotating) | $p_s$ | 5904.6 | 468.4 | 6017.0 | 598.0 |
| 2 | Mobile transects (rotating) | $p_I$ | 5934.5 | 463.1 | 6063.0 | 614.5 |
| 2 | Mobile transects (rotating) | $p_I, p_s, p_F$ | 4608.8 | 830.0 | 4618.0 | 1162.0 |
| 3 | Distribution of turning angles | RW | 0.2539 | 0.0061 | 0.2540 | 0.0083 |
| 3 | Distribution of turning angles | $p_F$ | 0.2537 | 0.0061 | 0.2535 | 0.0077 |
| 3 | Distribution of turning angles | $p_s$ | 0.2546 | 0.0074 | 0.2545 | 0.0098 |
| 3 | Distribution of turning angles | $p_I$ | 0.2542 | 0.0061 | 0.2540 | 0.0086 |
| 3 | Distribution of turning angles | $p_I, p_s, p_F$ | 0.2502 | 0.0070 | 0.2501 | 0.0095 |
| 3 | Home range (Kernel estimate) | RW | 1.53E+08 | 1.89E+07 | 1.52E+08 | 2.10E+07 |
| 3 | Home range (Kernel estimate) | $p_F$ | 1.52E+08 | 1.88E+07 | 1.51E+08 | 2.19E+07 |
| 3 | Home range (Kernel estimate) | $p_s$ | 1.05E+08 | 1.07E+07 | 1.05E+08 | 1.41E+07 |
| 3 | Home range (Kernel estimate) | $p_I$ | 1.53E+08 | 1.98E+07 | 1.51E+08 | 2.12E+07 |
| 3 | Home range (Kernel estimate) | $p_I, p_s, p_F$ | 1.01E+08 | 1.15E+07 | 1.02E+08 | 1.57E+07 |
| 3 | Dilation | RW | 3.29E+06 | 1.41E+06 | 3.35E+06 | 2.20E+06 |
| 3 | Dilation | $p_F$ | 3.23E+06 | 1.41E+06 | 3.15E+06 | 2.19E+06 |
| 3 | Dilation | $p_s$ | 3.27E+06 | 1.34E+06 | 3.39E+06 | 2.04E+06 |
| 3 | Dilation | $p_I$ | 3.41E+06 | 1.41E+06 | 3.45E+06 | 2.19E+06 |
| 3 | Dilation | $p_I, p_s, p_F$ | 3.31E+06 | 1.35E+06 | 3.29E+06 | 2.05E+06 |
| 3 | Immobile transects | RW | 8219.5 | 124.0 | 8238.5 | 142.0 |
| 3 | Immobile transects | $p_F$ | 8221.1 | 117.0 | 8232.5 | 146.0 |
| 3 | Immobile transects | $p_s$ | 8140.2 | 133.7 | 8163.5 | 166.0 |
| 3 | Immobile transects | $p_I$ | 8223.7 | 113.5 | 8238.5 | 140.5 |
| 3 | Immobile transects | $p_I, p_s, p_F$ | 8124.4 | 146.2 | 8148.5 | 165.5 |
| 3 | Mobile transects (linear) | RW | 3805.0 | 297.3 | 3844.0 | 391.0 |
| 3 | Mobile transects (linear) | $p_F$ | 3791.9 | 295.8 | 3814.0 | 398.0 |
| 3 | Mobile transects (linear) | $p_s$ | 3810.2 | 284.6 | 3839.5 | 377.5 |
| 3 | Mobile transects (linear) | $p_I$ | 3785.0 | 288.0 | 3826.0 | 392.5 |
| 3 | Mobile transects (linear) | $p_I, p_s, p_F$ | 3611.9 | 358.0 | 3665.5 | 491.0 |
| 3 | Mobile transects (rotating) | RW | 8521.5 | 559.2 | 8686.0 | 648.0 |
| 3 | Mobile transects (rotating) | $p_F$ | 8538.0 | 511.2 | 8655.0 | 686.0 |
| 3 | Mobile transects (rotating) | $p_s$ | 8511.4 | 540.2 | 8657.5 | 636.0 |
| 3 | Mobile transects (rotating) | $p_I$ | 8518.4 | 553.5 | 8687.0 | 627.0 |
| 3 | Mobile transects (rotating) | $p_I, p_s, p_F$ | 8095.0 | 815.8 | 8315.0 | 1057.5 |
| 4 | Distribution of turning angles | RW | 0.2440 | 0.0062 | 0.2438 | 0.0084 |
| 4 | Distribution of turning angles | $p_F$ | 0.2442 | 0.0061 | 0.2443 | 0.0078 |
| 4 | Distribution of turning angles | $p_s$ | 0.2444 | 0.0073 | 0.2444 | 0.0105 |
| 4 | Distribution of turning angles | $p_I$ | 0.2438 | 0.0060 | 0.2436 | 0.0082 |
| 4 | Distribution of turning angles | $p_I, p_s, p_F$ | 0.2406 | 0.0067 | 0.2407 | 0.0093 |
| 4 | Home range (Kernel estimate) | RW | 1.38E+08 | 1.99E+07 | 1.37E+08 | 2.20E+07 |
| 4 | Home range (Kernel estimate) | $p_F$ | 1.39E+08 | 2.07E+07 | 1.36E+08 | 2.12E+07 |
| 4 | Home range (Kernel estimate) | $p_s$ | 9.74E+07 | 1.09E+07 | 9.76E+07 | 1.48E+07 |
| 4 | Home range (Kernel estimate) | $p_I$ | 1.37E+08 | 1.94E+07 | 1.35E+08 | 2.17E+07 |
| 4 | Home range (Kernel estimate) | $p_I, p_s, p_F$ | 9.18E+07 | 1.25E+07 | 9.19E+07 | 1.67E+07 |
| 4 | Dilation | RW | 2.64E+06 | 1.37E+06 | 2.64E+06 | 2.17E+06 |
| 4 | Dilation | $p_F$ | 2.59E+06 | 1.35E+06 | 2.59E+06 | 2.00E+06 |
| 4 | Dilation | $p_s$ | 2.50E+06 | 1.31E+06 | 2.55E+06 | 2.04E+06 |
| 4 | Dilation | $p_I$ | 2.70E+06 | 1.39E+06 | 2.72E+06 | 2.11E+06 |
| 4 | Dilation | $p_I, p_s, p_F$ | 2.63E+06 | 1.22E+06 | 2.61E+06 | 1.86E+06 |
| 4 | Immobile transects | RW | 5369.7 | 313.3 | 5402.0 | 419.0 |
| 4 | Immobile transects | $p_F$ | 5370.1 | 297.2 | 5405.0 | 385.0 |
| 4 | Immobile transects | $p_s$ | 5523.5 | 313.2 | 5547.5 | 431.0 |
| 4 | Immobile transects | $p_I$ | 5363.3 | 308.1 | 5396.0 | 394.0 |
| 4 | Immobile transects | $p_I, p_s, p_F$ | 5340.8 | 366.2 | 5387.5 | 506.0 |
| 4 | Mobile transects (linear) | RW | 2999.6 | 326.7 | 3033.5 | 429.5 |
| 4 | Mobile transects (linear) | $p_F$ | 2998.7 | 317.6 | 3033.0 | 419.0 |
| 4 | Mobile transects (linear) | $p_s$ | 3004.7 | 306.8 | 3020.5 | 420.5 |
| 4 | Mobile transects (linear) | $p_I$ | 2973.5 | 314.5 | 3010.0 | 421.5 |
| 4 | Mobile transects (linear) | $p_I, p_s, p_F$ | 2720.3 | 393.4 | 2761.0 | 532.0 |
| 4 | Mobile transects (rotating) | RW | 7056.5 | 584.9 | 7234.0 | 683.5 |
| 4 | Mobile transects (rotating) | $p_F$ | 7077.2 | 590.2 | 7257.0 | 689.0 |
| 4 | Mobile transects (rotating) | $p_s$ | 7021.6 | 578.3 | 7176.0 | 728.5 |
| 4 | Mobile transects (rotating) | $p_I$ | 7032.4 | 569.2 | 7199.0 | 710.0 |
| 4 | Mobile transects (rotating) | $p_I, p_s, p_F$ | 6385.6 | 918.6 | 6543.0 | 1346.5 |
| 5 | Distribution of turning angles | RW | 0.2440 | 0.0067 | 0.2440 | 0.0093 |
| 5 | Distribution of turning angles | $p_F$ | 0.2444 | 0.0065 | 0.2445 | 0.0088 |
| 5 | Distribution of turning angles | $p_s$ | 0.2442 | 0.0077 | 0.2442 | 0.0104 |
| 5 | Distribution of turning angles | $p_I$ | 0.2445 | 0.0064 | 0.2443 | 0.0089 |
| 5 | Distribution of turning angles | $p_I, p_s, p_F$ | 0.2458 | 0.0075 | 0.2456 | 0.0104 |
| 5 | Home range (Kernel estimate) | RW | 1.43E+08 | 1.89E+07 | 1.42E+08 | 2.20E+07 |
| 5 | Home range (Kernel estimate) | $p_F$ | 1.44E+08 | 1.92E+07 | 1.43E+08 | 1.97E+07 |
| 5 | Home range (Kernel estimate) | $p_s$ | 9.76E+07 | 9.84E+06 | 9.79E+07 | 1.28E+07 |
| 5 | Home range (Kernel estimate) | $p_I$ | 1.43E+08 | 1.77E+07 | 1.42E+08 | 2.04E+07 |
| 5 | Home range (Kernel estimate) | $p_I, p_s, p_F$ | 8.35E+07 | 1.74E+07 | 8.03E+07 | 2.23E+07 |
| 5 | Dilation | RW | 4.08E+06 | 1.38E+06 | 4.17E+06 | 2.09E+06 |
| 5 | Dilation | $p_F$ | 4.07E+06 | 1.40E+06 | 4.07E+06 | 2.10E+06 |
| 5 | Dilation | $p_s$ | 3.85E+06 | 1.35E+06 | 3.86E+06 | 2.03E+06 |
| 5 | Dilation | $p_I$ | 4.16E+06 | 1.40E+06 | 4.17E+06 | 2.15E+06 |
| 5 | Dilation | $p_I, p_s, p_F$ | 3.91E+06 | 1.11E+06 | 3.90E+06 | 1.57E+06 |
| 5 | Immobile transects | RW | 7113.1 | 174.9 | 7133.0 | 217.5 |
| 5 | Immobile transects | $p_F$ | 7118.3 | 172.7 | 7141.5 | 211.0 |
| 5 | Immobile transects | $p_s$ | 7110.6 | 181.6 | 7141.0 | 229.5 |
| 5 | Immobile transects | $p_I$ | 7117.0 | 178.7 | 7146.5 | 217.0 |
| 5 | Immobile transects | $p_I, p_s, p_F$ | 6566.2 | 454.3 | 6623.5 | 590.0 |
| 5 | Mobile transects (linear) | RW | 2973.3 | 284.4 | 3005.0 | 381.0 |
| 5 | Mobile transects (linear) | $p_F$ | 2978.3 | 290.1 | 3000.5 | 390.5 |
| 5 | Mobile transects (linear) | $p_s$ | 2992.5 | 275.5 | 3020.0 | 346.5 |
| 5 | Mobile transects (linear) | $p_I$ | 2988.9 | 281.2 | 3013.0 | 387.5 |
| 5 | Mobile transects (linear) | $p_I, p_s, p_F$ | 1907.4 | 443.2 | 1905.0 | 634.0 |
| 5 | Mobile transects (rotating) | RW | 6987.0 | 496.1 | 7126.0 | 563.5 |
| 5 | Mobile transects (rotating) | $p_F$ | 6963.1 | 505.5 | 7100.5 | 628.0 |
| 5 | Mobile transects (rotating) | $p_s$ | 6964.0 | 500.3 | 7105.0 | 611.0 |
| 5 | Mobile transects (rotating) | $p_I$ | 7015.8 | 448.2 | 7142.0 | 523.5 |
| 5 | Mobile transects (rotating) | $p_I, p_s, p_F$ | 4427.2 | 1074.3 | 4417.5 | 1546.0 |
